# Mass mortality associated with Carpione Rhabdovirus in golden pompano (*Trachinotus ovatus*) in China: First report

**DOI:** 10.1101/2024.05.14.594202

**Authors:** Heng Sun, Jie Huang, Haoyu Wang, Yule Zhang, Qing Fei, Jie Zhou, Lindi Yang, Yanping Li, Shuanghu Cai, Yucong Huang

**Affiliations:** Fisheries College of Guangdong Ocean University, Guangdong Provincial Key Laboratory of Aquatic Animal Disease Control and Healthy Culture & Key Laboratory of Control for Diseases of Aquatic Economic Animals of Guangdong Higher Education Institutes, Zhanjiang, China

**Keywords:** CARPV, *Trachinotus ovatus*, Virus detection and isolation, Initial infection, Next-generation sequencing

## Abstract

A severe epizootic outbreak occurred in a deep-water cultured golden pompano (*Trachinotus ovatus*) in Guangdong Province, China (August–November 2023); the cumulative mortality was 65–82%. The diseased fish showed clinical signs of lethargy, anorexia, whirling movements, and hemorrhage at the base of the fins and in the upper and lower jaws before mass mortality. A Rhabdovirus strain, CARPV2023, was successfully isolated using FHM cells. Cytopathic effects of the tissue filtrate of the diseased golden pompano on FHM and EPC cell monolayers were characterized by rounded cells, grape-like cluster formation, detachment, and lysis. Histopathology revealed congestion, hemorrhage, and necrosis in the spleen, liver, kidney, and intestinal tissues of naturally and experimentally infected golden pompanos. Electron microscopy showed that bullet-shaped viral particles 183.5–201.6-nm long and 57.3–82.7-nm wide budded from cell membrane or aggregated around the infected FHM cells. The virus remained stable for 7 days at 4–33°C and grew optimally at 28°C. Whole-genome sequencing and phylogenetic analysis based on the full genome further indicated that the CARPV2023 strain is a member of Carpione Rhabdovirus, which is related to *Novirhabdovirus* with unclassification. A specific nested reverse transcriptase-polymerase chain reaction confirmed the presence of CARPV in the affected golden pompano. Much higher mortality was observed in challenged golden pompanos than in the controls through intraperitoneal injection and immersion infection. In situ hybridization showed positive reactions in the fish tissues. This is the first study to report natural CARPV infections in marine fish in the world.

## Introduction

Rhabdoviruses are a class of negative-sense viruses with single-stranded RNA that can infect a range of hosts, including mammals, birds, reptiles, fish, insects, and plants [1,2]. According to the latest viral taxonomy profile published by the International Committee on Taxonomy of Viruses (ICTV) in 2022, Rhabdoviridae includes three subfamilies, 46 genera, and 318 species of viruses [3]. Virions are typically bullet- or rod-shaped and are enveloped by an outer membrane; however, they can also be filamentous without an envelope. The viral genome is approximately 10–16 kb long and encodes at least five structural proteins: nucleoprotein (N), phosphoprotein (P), matrix protein (M), glycoprotein (G), and RNA-dependent RNA-Polymerase (L). The five viral genes are organized in the order type of rhabdoviruses: 3′-N–P–M–G–L-5′[1,4]. Nevertheless, additional genes encoding other proteins or alternative open reading frames (ORFs) have been found in the genome. Among these, fish rhabdoviruses are widely distributed in marine and freshwater fish worldwide and cause serious diseases and huge economic losses [3]. To date, more than 20 rhabdoviruses have been reported in fish, including the infectious hematopoietic necrosis virus (IHNV), viral hemorrhagic septicemia virus (VHSV), spring viremia of carp virus (SVCV), snakehead rhabdovirus (SHRV), Siniperca chuatsi rhabdovirus (SCRV), perch rhabdovirus (PRV), and Pike fry rhabdovirus (PFRV) [5,6]. Rhabdoviruses originating from fish mainly belong to five genera: *Novirhadovirus*, *Perhabdovirus*, *Siniperhavirus*, *Sprivivirus*, and *Scophrhavirus*; however, some have not yet been assigned [3–7].

The golden pompano (*Trachinotus ovatus*), an economically important marine fish species in the genus *Trachinotus* of the family Carangidae, is widely distributed in the subtropical and tropical sea areas of Southeast Asian countries. In recent years, the golden pompano has been cultured on a large scale in offshore cages along the southern coast of China, owing to its fast growth rate, strong environmental adaptability, and remarkable economic value. According to the Fishery Statistical Yearbook of China, the annual production of golden pompano in China will reach 243,908 tons by 2022 [8].

However, owing to the germplasm degradation, excessive density, unbalanced nutrition, and decline in cultural environment, the cultural golden pompano has frequently suffered from infectious diseases caused by *Cryptocaryon irritans*, *Nocardia seriolae*, *Photobacterium damselae*, and *Streptococcus* sp. [9–17]. Diseases have become a key constraint to the healthy and sustainable development of golden pompano aquaculture. Recently, a novel epidemic occurred and caused mass mortality in the golden pompano in China. The present study describes the clinical signs, histological characteristics, viral culture, whole-genome sequencing analysis of viruses, physicochemical properties of viruses, reverse transcriptase-polymerase chain reaction (RT-PCR) detection, and infection trials in farmed golden pompanos in China.

## Materials and Methods

### Cases history and fish sampling

From August–November 2023, an acute infectious disease occurred in golden pompanos cultured in offshore cages in Guangdong Province, China. The affected fish were housed at a stocking density of 80,000–100,000 fish per cage, with a circumference of 80–90 m and a depth of 9 m. The onset of disease and mortality was observed in August 2023, approximately 4 months after the golden pompano fry were introduced into the cages. Approximately 300–2000 fish deaths in each cage were recorded daily, with mortality peaking 5–9 days after the first noticeable mortality. The water temperature of affected cages during the epidemic period was 24.2–30.7°C, and dissolved oxygen concentration was 4.85–7.76 mg/L. Ammonia and nitrate concentrations were 0.063–0.186 and 0.011–0.024 mg/L, respectively. Samples of moribund golden pompanos weighing 183–455 g were randomly collected from five offshore cage farms and immediately sent to the laboratory in aerated water bags for viral isolation, histopathological examination, and RT-PCR detection.

### Parasitological and Microbiological Examination

The exterior mucus, gills, and viscera of the clinically diseased *T. ovatus* samples were examined using a light microscope (ZEISS, DM750, Germany) according to the methods described by Huang et al. [17]. For bacterial isolation, brain, spleen, and kidney tissue from moribund fish were inoculated onto Zobell marine agar 2216E and incubated on brain heart infusion agar (BHIA) (Hopebio, Qingdao, China) at 28°C for 2 days. A single colony was selected for further isolation and purification, followed by expansion in the BHI broth. Genomic DNA was extracted from the bacterial suspension using a DNA extraction kit (Tiangen, China) and subsequently subjected to PCR using universal 16s rRNA primers [18]. The amplified products were sent to the Sangon Company (China) for sequencing. The obtained sequencing data were compared with those hosted on the NCBI database to identify the causative bacterial species. The nervous necrosis virus (NNV), megalocytivirus, infectious hematopoietic necrosis virus (IHNV), and viral hemorrhagic septicemia virus (VHSV) were detected using PCR or RT-PCR to screen for the presence of these viruses in fish samples, according to the protocol described by the OIE standard and Huang et al. (Table1) [17,19–24].

### Cells

EPC cells were grown in Minimum Essential Medium-α (MEM-α, Gibco) supplemented with 10% fetal bovine serum (FBS, Sangon Co Ltd, Shanghai, China) at 28 °C in a humidified 5% CO2 incubator. FHM[25] cells were grown in Leibovitz’s L-15 (L-15, Biosharp) supplemented with 10% FBS at 28 °C.

### Viral Culture

The tissue samples from the spleen and kidney of affected golden pompano were dissected and homogenized using a MICCRA D-1 homogenizer (Miccra Gmbh, Müllheim, Germany) in Leibovitz L-15 medium containing penicillin (100 IU/mL), streptomycin (100 mg/mL), and kanamycin (100 mg/mL). The homogenates were then centrifuged at 3,000 g at 4 °C for 10 min, followed by filtration of the supernatant twice through a 0.22 μm filter membrane. Subsequently, the supernatants were inoculated at dilutions of 1:10 and 1:100 onto FHM cells grown in 25 cm^2^ flasks containing Leibovitz medium (L-15) supplemented with 2% FBS, penicillin (100 IU/mL), and streptomycin (100 mg/mL) at 25 °C. The inoculated cells were monitored daily for cytopathic effects (CPEs) under a light microscope (DMi8, Leica, Germany). The supernatants of the cultures that developed CPEs were sampled and processed for RT-PCR as described above.

### Electron microscopy

Viral stock solutions were separately inoculated into the FHM monolayer cells. After the development of cytopathic effects, the cells were harvested and centrifuged at 1,000 ×*g* for 10 min. Precipitation was fixed in 2.5% glutaraldehyde in phosphate-buffered saline and post-fixed in 1% osmium tetroxide. Subsequently, fixed cells were subjected to gradient dehydration in ethanol and embedded in 618 epoxy resin (Leica Ultracul-R). Ultrathin sections were prepared using a microtome (Leica, EM UC7), double-stained with uranyl acetate and lead citrate, and examined under a JEM1230 transmission electron microscope (Nippon Electronics, Japan).

### Next-generation sequencing of virus

Viral DNA was extracted directly from 200 µL aliquots of infected cell culture and nuclease treated using a commercial QIAamp DNA Kit (Qiagen) according to the manufacturer’s procedures. RNA was extracted from samples of infected cell lysates using TRIzol reagent and reverse transcribed using random hexamers, followed by second-strand synthesis. Library construction and Illumina sequencing were performed by Tpbio Co., Ltd. (Shanghai, China). Paired-end 150 nt reads were generated using the Illumina NovaSeq 6000 System (Illumina, San Diego, CA, USA).

### Genome assembling and phylogenetic analysis

Raw reads were filtered and trimmed using Fastp (https://github.com/OpenGene/fastp) to remove the sequencing adapters and low-quality reads. Ribosomal RNAs and host read subtraction by read mapping were performed using the BBMAP program. De novo genome assembly was performed using SPAdes v3.13.0. Single nucleotide polymorphisms (SNPs) were identified using the integrated software snippy (v4.4.5), which included both substitutions (SNPs) and insertions/deletions (indels). The phylogenetic tree was constructed based on the complete genome sequences of the isolated virus from golden pompano and other Rhabdomyviridae viruses retrieved from GenBank using MEGA X software with the maximum likelihood method and 1000 bootstrap replicates.

### Biological characteristics of Virus

#### Viral titer determination

The EPC and FHM cells were transferred into 96-well plates and incubated at 25 °C. The viral isolate was serially diluted in the cell culture medium [26]. Each well of a 96-well plate received 100 µL of the diluted virus suspension. The cells were then incubated in a controlled environment at 28 °C until no additional CPEs could be observed. Following formaldehyde fixation, the inoculated cells were stained using crystal violet solution, and the TCID_50_ values were calculated using the Reed and Muench method (1938) [27].

#### Viral infection at different temperatures

To determine the effect of temperature on the isolated virus infection, FHM cells were incubated with CAPRV2023 (TCID_50_=10^7.72^/0.1 mL) at 28°C for 1 h with agitation every 10 min. Subsequently, the virus was removed and replaced with 5 mL of L-15 medium containing 2% FBS. Finally, the infected cell culture flasks were placed in separate incubators set at 23, 28, and 33 °C.

#### Thermal and UV inactivation of virus

The isolated virus suspension was placed separately in water baths set to 50 and 55°C. Samples were collected every 5 min to determine the duration of inactivation. To investigate the inactivation time of CAPRV2023 under UV light, L-15 medium (2% FBS) containing TCID_50_=10^6.72^/0.1 mL was placed in cell culture dishes within a biosafety chamber (MSC Advantage 1.5, Thermo scientific, USA) and exposed to UV radiation from a lamp emitting at 254 nm. The radiation intensity was set to 400 mW/m^2^. The samples were collected every 15 s for an irradiation period of 60 s. Viral titers were measured immediately after each irradiation interval.

#### Survives of virus in natural seawater

To determine the survival of the virus isolated in nature, 1 L of natural seawater was obtained from ocean near the fish farm, sterilized at 121 °C for 15 min, and cooled at room temperature for 2 h. After natural precipitation of inactivated plankton and other particles in the seawater, the supernatant was transferred to a 50 mL centrifuge tube and centrifuged at 12000 rpm for 10 min and collected for further use. The viral suspension (TCID_50_= 10^6.72^/0.1 mL) diluted with natural seawater was aliquoted into a cryopreservation tube (1 mL per tube). The samples were stored at 28 °C for f 1, 3, 7, 14, 21, and 28 days. Subsequently, the TCID_50_ of the viral dilution was determined as described above.

#### Growth kinetics of virus

The virus was diluted to a concentration of TCID_50_=10^3.72^/0.1 mL in L-15 medium supplemented with 10% FBS. Subsequently, it was inoculated onto FHM cells and incubated at 23, 28, and 33°C for 12, 24, 36, 48, and 72 h. Following lysis, the supernatants were collected to determine the TCID_50_.

### Reverse transcriptase-polymerase chain reaction

For RNA extraction from total RNA, approximately 50 mg spleen, kidney, and brain tissues from diseased golden pompanos and infected cell cultures were extracted using a commercial RNA extraction kit for marine animals (Tiangen, Beijing, China). The concentration and purity of the extracted RNA were determined using Nanodrop 2000 (Thermo Scientific, Waltham, MA, USA). Total RNA was reverse transcribed into complementary DNA (cDNA) using random primers and M-MLV reverse transcriptase (TransGen Biotech, Beijing, China). To detect the presence of viral pathogens in cultured golden pompano and infected cell cultures, primers for nested PCR were designed based on G protein gene sequences after sequencing the virus isolated from diseased golden pompano (Table 1). For the first round RT-PCR amplification, the cDNA was used as template with the primers CAPRV2023-F1/R1 at an annealing temperature of 55°C. For the nested PCR, the product of the first round PCR was used as the template with the primers CAPRV2023-F2/R2 at an annealing temperature of 59°C. DNA fragments of 802 bp and 190 bp were amplified using the first- and second-round RT-PCR, respectively. The PCR products were purified using a PCR purification kit (Tiangen, Beijing, China) and sequenced by Sangon Biotech Co., Ltd. (Shanghai, China).

**Table 1.**
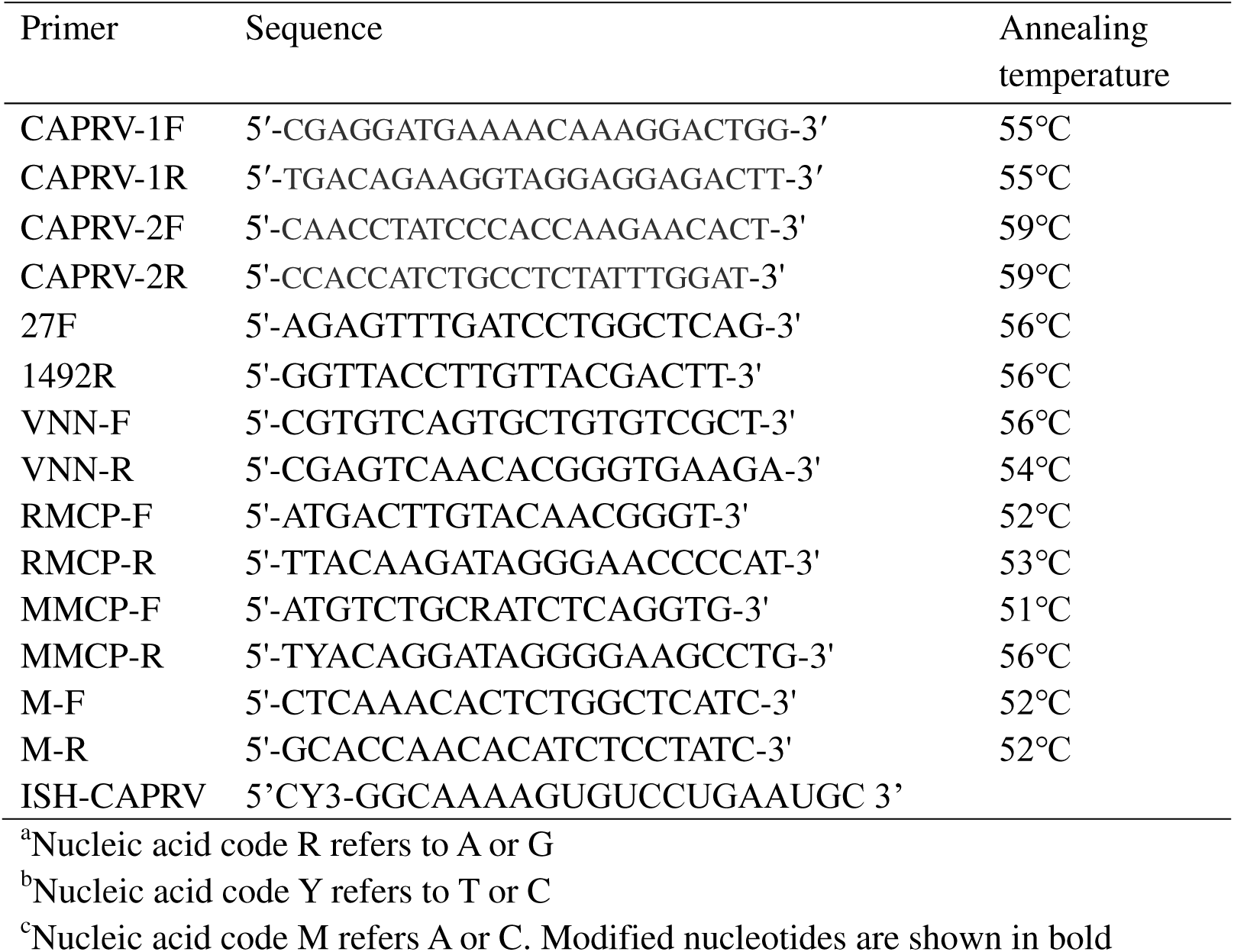
Primer sequences used in this study.

**Table 2.** Titers of CAPRV with different treatments.

| Treatment method | TCID<sub>50</sub> ml<sup>-1</sup> |
| --- | --- |
| Control group | 10<sup>7.33</sup> |
| 55℃ (10min) | 0 |
| 50℃ (15min) | 10<sup>1.68</sup> |
| 50℃ (18min) | 0 |
| 4℃ (40days) | 10<sup>2.88</sup> |

### Histopathology

The liver, spleen, kidney, heart, intestine, and brain tissues from the naturally infected golden pompano*s* were fixed in 4% neutral paraformaldehyde fixative (Sigma, USA) for 24 h, followed by alcohol gradient dehydration, xylene transparency before being embedded in paraffin wax according to standard procedures. The samples were then cut into 4–6 μm thick sections with a microtome (RM2245, Leica, Germany). The sections were stained with hematoxylin and eosin (HE) and examined under a light microscope equipped with a CCD imaging system (Axio Scope 5, Zeiss, Germany).

### Fluorescence in situ hybridization

To investigate tissue-specific infection of the isolated virus in golden pompano, a probe was designed for fluorescence in situ hybridization based on the gene sequence of the CAPRV G protein (Table 1). The tissue sections were treated with 0.2M HCl for 5 min at room temperature, followed by three washes with PBS for 5 min each. Subsequently, the sample were incubated with proteinase K (40 μg/mL) at room temperature for 20 min. The samples were then washed again three times with PBS containing 0.2% glycine for 5 min each. After fixation with 4% paraformaldehyde (PFA, Sigma, USA) for 10 min and two additional washes with PBS for 5 min each, the samples were incubated with a solution of acetic anhydride and triethanolamine at room temperature for 10 min and washed twice with PBS for 5 min each. Next, the prehybridization reaction was carried out at 44°C for 2 h, followed by overnight hybridization at 44°C. Subsequently, the samples underwent two washes with saline-sodium citrate (SSC) buffer five times consecutively, where each wash lasted for 20 min. The sample were then subjected to three washes in a mixture of deionized formamide (50%) and SSC (2X) at 37°C for 20-min intervals before being washed five times in Tris-buffered saline Tween-20 (TBST) lasting 5 min per wash. Finally, the slides were counterstained with DAPI and briefly washed with TBST before being sealed with a fluorescent inhibitor to facilitate microscopic examination and photography (LSM880; ZEISS, Germany).

### Experimental infection

A total to 600 healthy golden pompano (body weight, 75–92 g) were purchased from a fish farm with no history of disease and acclimatized to 8 m^3^ well-aerated cement ponds filled with clean sea water at 28–30°C for 1 week. The fish were fed twice daily with commercial fish feed and confirmed to be pathogen-free prior to experimental infection. The fish were randomly divided into six replicate groups (20 fish per replicate). Fish from the tissue homogenate injection group and the cell supernatant injection group were injected intraperitoneally (i.p.) with 0.2 mL tissue homogenate, as described above, and 10^5^ TCID_50_/fish of virus, respectively. Correspondingly, fish from the two control groups were intraperitoneally injected with the same dose of 1:10 diluted normal FHM cell culture supernatant and PBS. Fish from the tissue homogenate immersion group were exposed to TCID_50_ (10^5^/0.1 mL of CARPV2023 by immersion for 2 h, and fish from the control group were immersed in 1:100 diluted FHM cell culture supernatants. After infection, all fish were transferred to the corresponding cement ponds. Throughout the study period, all fish were maintained in well-aerated pools at a temperature of 28°C. Clinical signs and mortality were monitored daily, and dead or moribund fish from each group were collected for PCR, as described above, to determine the presence of pathogens.

## Results

### Disease characterization, clinical signs, and autopsy analysis

High mortalities were experienced and the estimated cumulative mortality among golden pompano farms during this outbreak was approximately 65–82% based on data obtained from cage farms during epidemiological investigations (Figure 1A). The diseased fish showed clinical signs of lethargy, anorexia, whirling movements in the water, and hemorrhage at the base of the fins and in the upper and lower jaws (Figure 1B). Autopsy revealed pale gills, abundant red ascites, hepatomegaly, splenomegaly, congestion, and hemorrhages in the liver, spleen, kidney, and intestine (Figure 1C).

**Figure 1.**
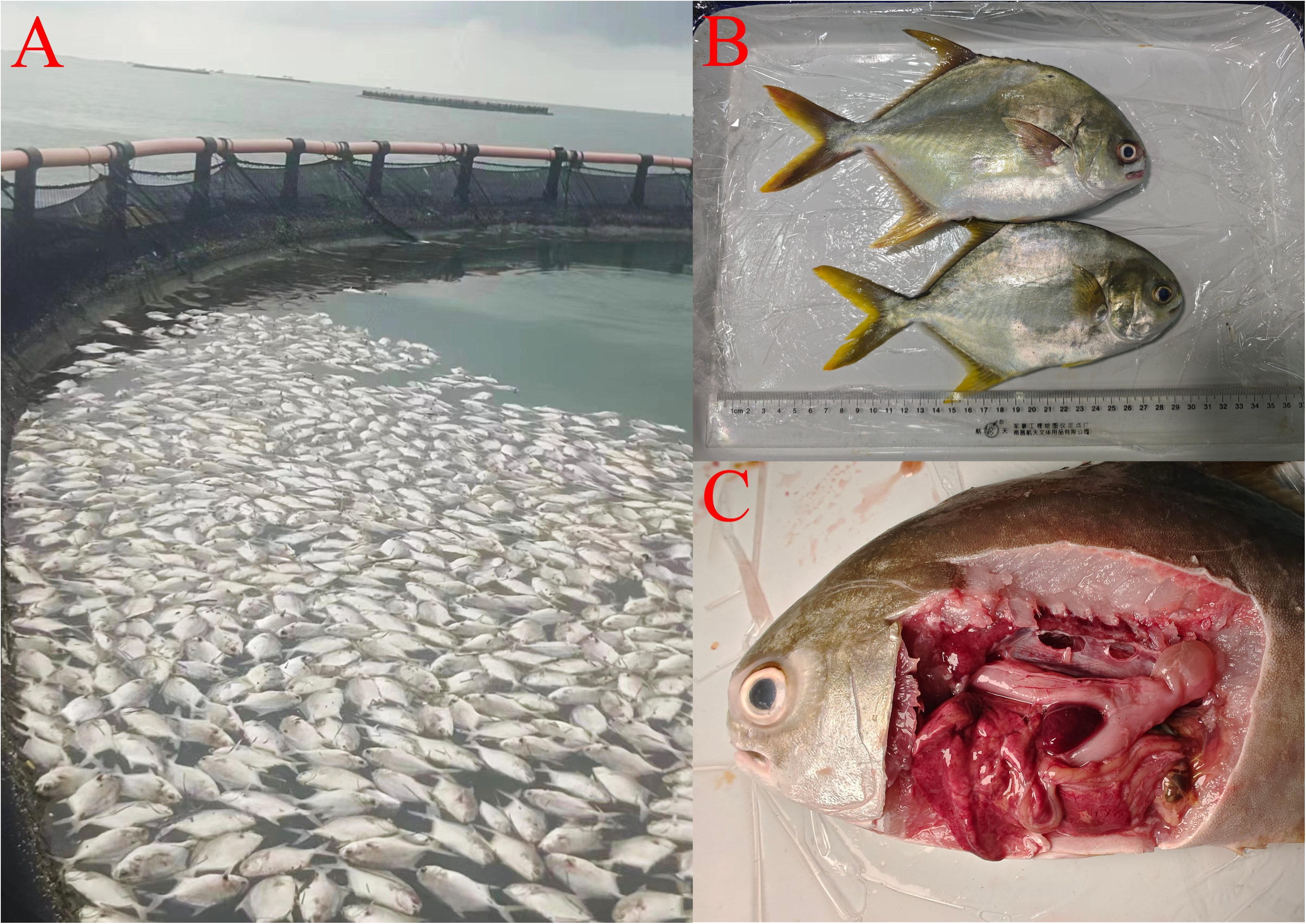
Golden pompano infected with CAPRV2023. (A) Significant mortality occurred in golden pompano cultured in off-shore cage, (B) naturally diseased golden pompano showing hemorrhages at the base of the fins and on the upper and lower jaws. (C) A naturally diseased golden pompano displaying pale gills, accumulation of red ascites, hepatomegaly, and splenomegaly and congested liver, spleen, kidney, and intestine.

### Isolation of virus

After 12 h of inoculation with the filtrate of tissues from the diseased fish, FHM and EPC cells developed classical CPEs consisting of rounded cells, grape-like cluster formation (Figure 2), and degeneration and lysis in the late stage. Following two to four passages of supernatants into fresh cell cultures, similar CPEs were observed.

**Figure 2.**
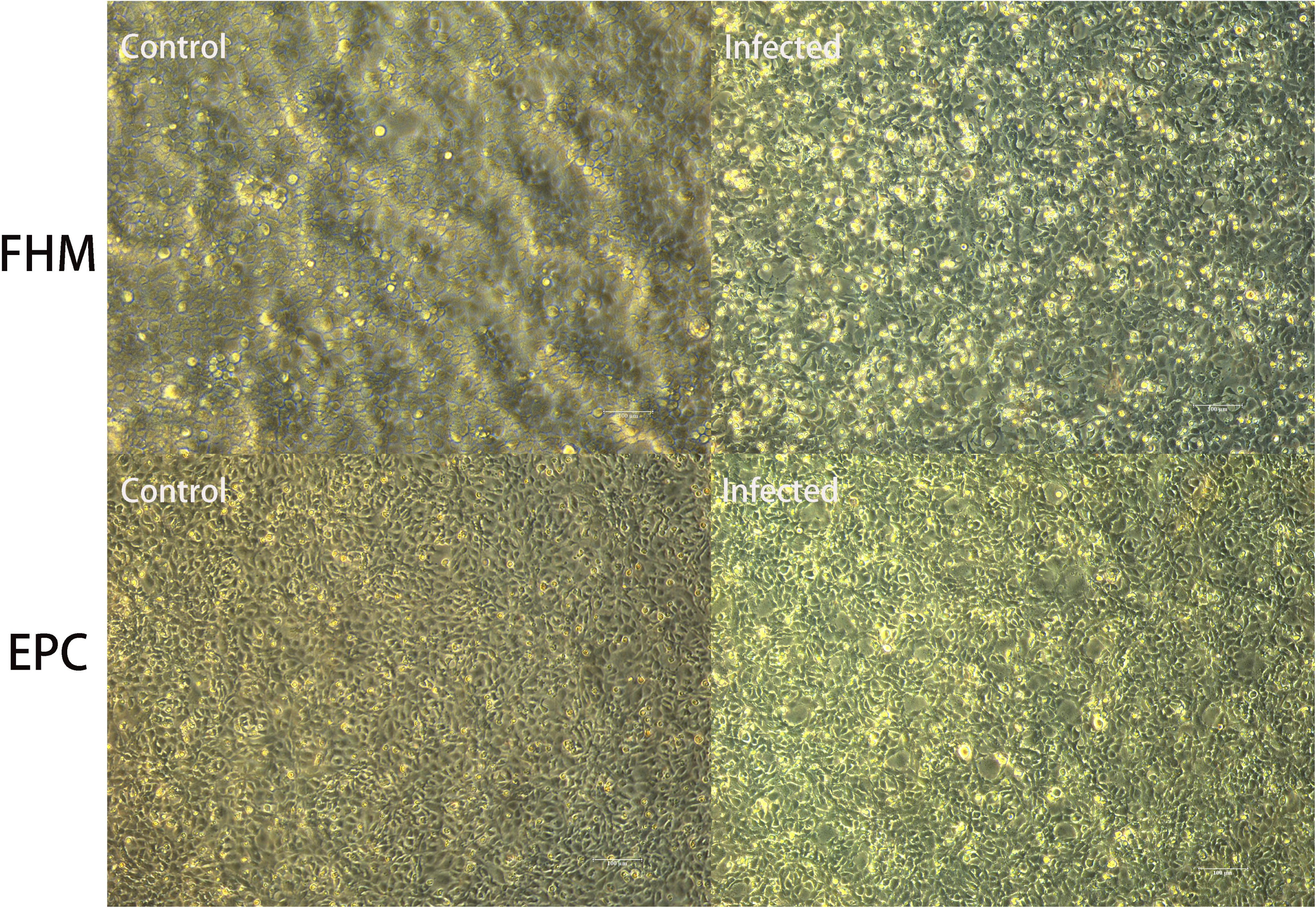
Cytopathic effects in FHM and EPC cells inoculated with tissue homogeneity filtrate from diseased golden pompano.

Transmission electron microscopy revealed that bullet-shaped virus particles budded or aggregated on the surface of the membrane in FHM cells infected with the viral isolate CAPRV2023, and the presence of CAPRV2023 encapsulated in vesicles could also be observed within the lysed cellular debris. (Figure 3A-B). Virus particles were also found within the cytoplasm of infected FHM cells (Figure 3C-D). Upon measurement, these virus particles were determined to be approximately 183.5–217.6 nm in length and 57.3–82.7 nm in width. The presence of CAPRV2023 encapsulated in vesicles was also observed in the lysed cellular debris.

**Figure 3.**
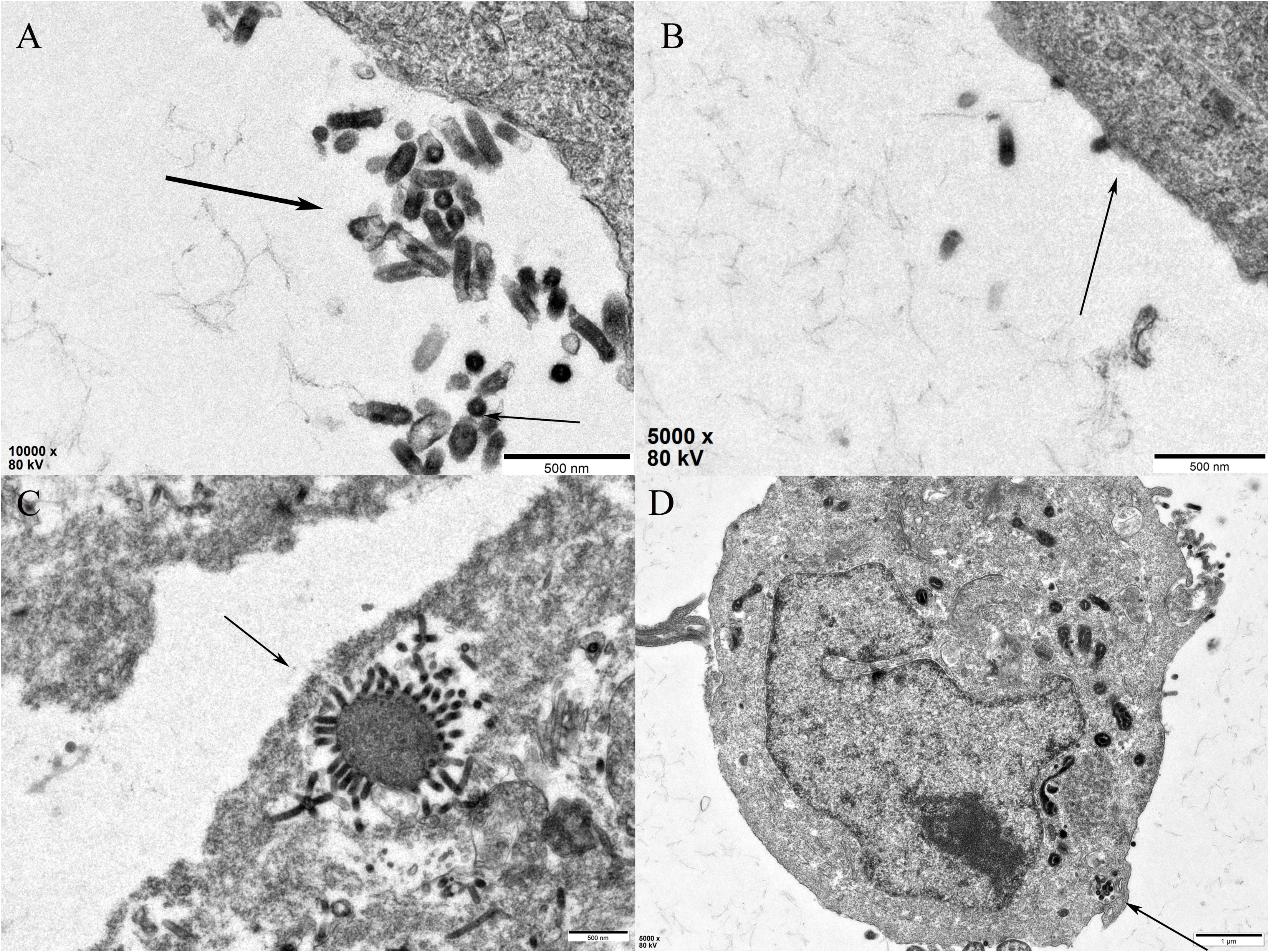
Transmission electron micro graphs of FHM cells infected with CAPRV2023. (A) viral particles of CAPRV2023 around the cells (indicated by black arrows), (B) CAPRV2023 budding from the surface of membrane. (C) CAPRV2023 encapsulated in vesicles observed in lysed cells (D) CAPRV2023 aggregating on the surface of membrane.

### Detection of virus

A nested RT-PCR assay was successfully established for highly sensitive detection of CAPRV. No DNA fragments were amplified in the first and second rounds of PCR using IHNV, VHSV, *Megalocytivirus*, *Ranavirus*, NNV, *S. iniae*, *S. dysgalactiae*, *S. agalactiae*, *L. garvieae*, *V. harveyi*, *V. alginolyticus*, and genomic DNA from healthy golden pompano as templates (Figure 4). Only 802 bp amplicon in the first-step PCR and 190 bp amplicon in the second-step PCR were obtained, indicating that the nested PCR was highly specific for CAPRV. All the tested golden pompanos collected from the four distinct sampling sites were negative for NNV, *Megalocytivirus*, IHNV, and VHSV, as revealed by PCR and RT-PCR (data not shown). Specific PCR products of 190 bp targeting the G protein gene were successfully detected in naturally and experimentally infected fish (Figure 5A) and in cultured FHM inoculated with the tissue filtrate of diseased fish by nested RT-PCR. Moreover, the distribution of CAPRV in the tissues of golden pompano was detected using RT-PCR with CAPRV-F2/CAPRV-R2 primers designed here, and the results showed that CAPRV was distributed in all organs examined (Figure 5B).

**Figure 4.**
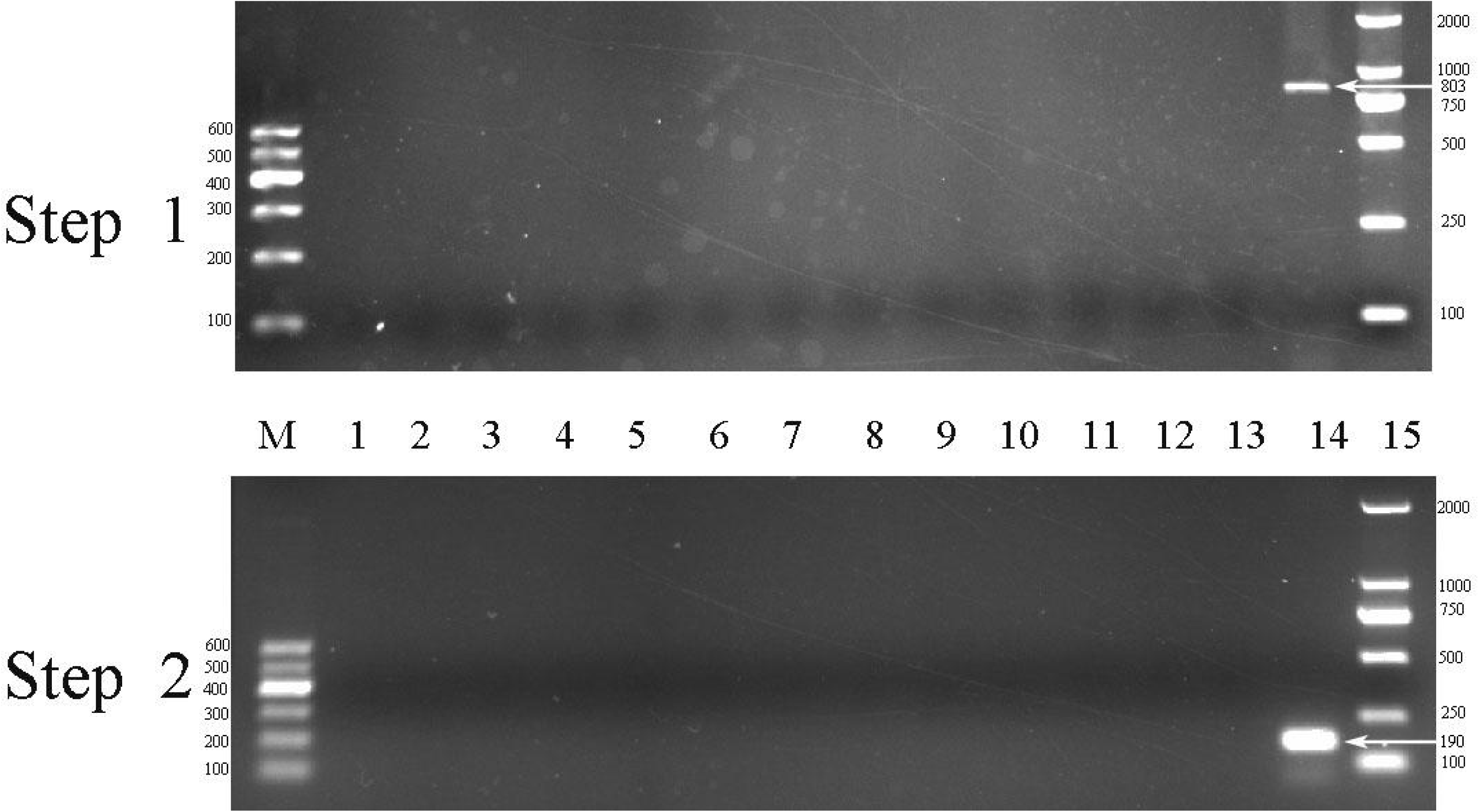
Specificity test with nested primers. Lane M: 600 bp DNA Marker, Lane 1: negative control, Lanes 2–14: IHNV, VHSV, *Megalocytivirus, Ranavirus*, nervous necrosis virus, *Streptococcus iniae*, *S. dysgalactiae*, *S. agalactiae*, *L. garvieae*, *V. harveyi*, *V. alginolyticus* and Genomic DNA from healthy golden pompano, Lane 15: 2000 bp DNA marker.

**Figure 5.**
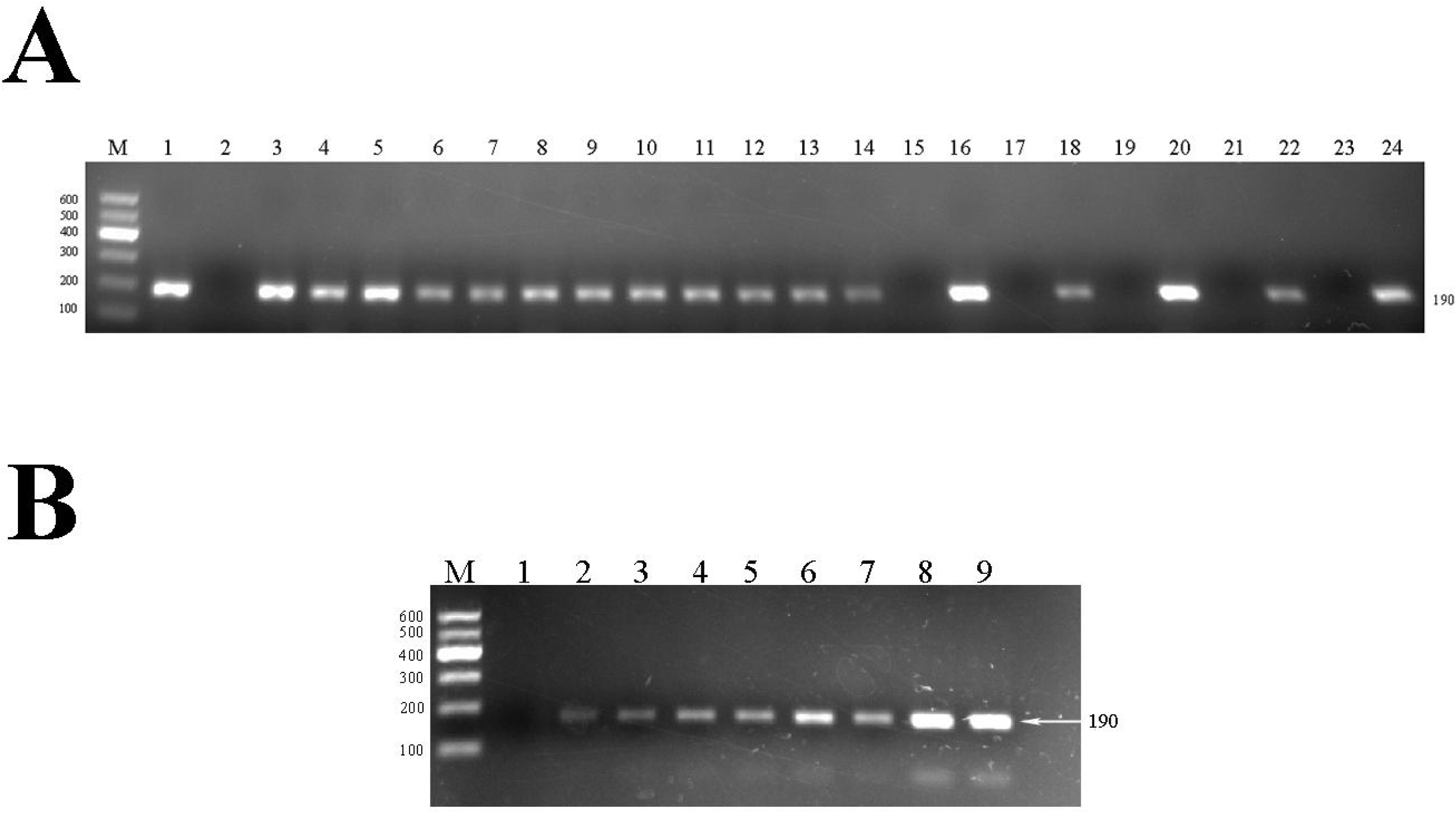
PCR test conducted on tissue samples. (A) Lane 1: negative control, Lanes 2– 9: brain, gill, heart, intestine, kidney, liver, spleen, and muscle. (B) Lane M: 600 DNA markers, Lane 1: positive control, Lane 2: negative control, Lanes 3–12: Naturally diseased golden pompano infected with CAPRV2023 from four sampling points, Lane15: fish from phosphate buffered saline injection group, Lane 16: fish from tissue homogenate injection group, Lane 17: fish from cell supernatant injection control group, Lane 18: fish from cell supernatant injection group, Lane 19: fish from immersion control group; Lane 20: fish from immersion group, Lane 21: healthy FHM cells, Lane 22: infected FHM cells, Lane 23: healthy EPC cells, Lane 24: infected EPC cells.

### Histology

Significant histopathological changes were detected in infected golden pompanos from different sampling sites, and the lesions were largely similar. Hepatocyte swelling and steatosis along with congestion in the hepatic sinuses and blood vessels were observed in the liver (Figure 6A). Splenic tissues showed focal necrosis in the splenic pulp, congestion, and hemorrhage in the splenic sinuses and blood vessels, accompanied by the deposition of melanin-macrophage centers (Figure 6B). In the kidney, degeneration, necrosis of renal tubule epithelial cells, focal necrosis, a large amount of inflammatory cell infiltration, and the presence of melanin-macrophage centers in the renal hematopoietic tissue were observed (Figure 6C). The epicardial layer was thickened, with inflammatory cell infiltration, indicating epicarditis (Figure 6D). The intestine displayed severe necrosis and exfoliation of the intestinal mucosal epithelial cells, a large amount of inflammatory cell infiltration in the mucosal layer, and dilated and congested blood vessels in the membrane propria and submucosa (Figure 6E). The brain showed notable cerebrovascular congestion (Figure 6F).

**Figure 6.**
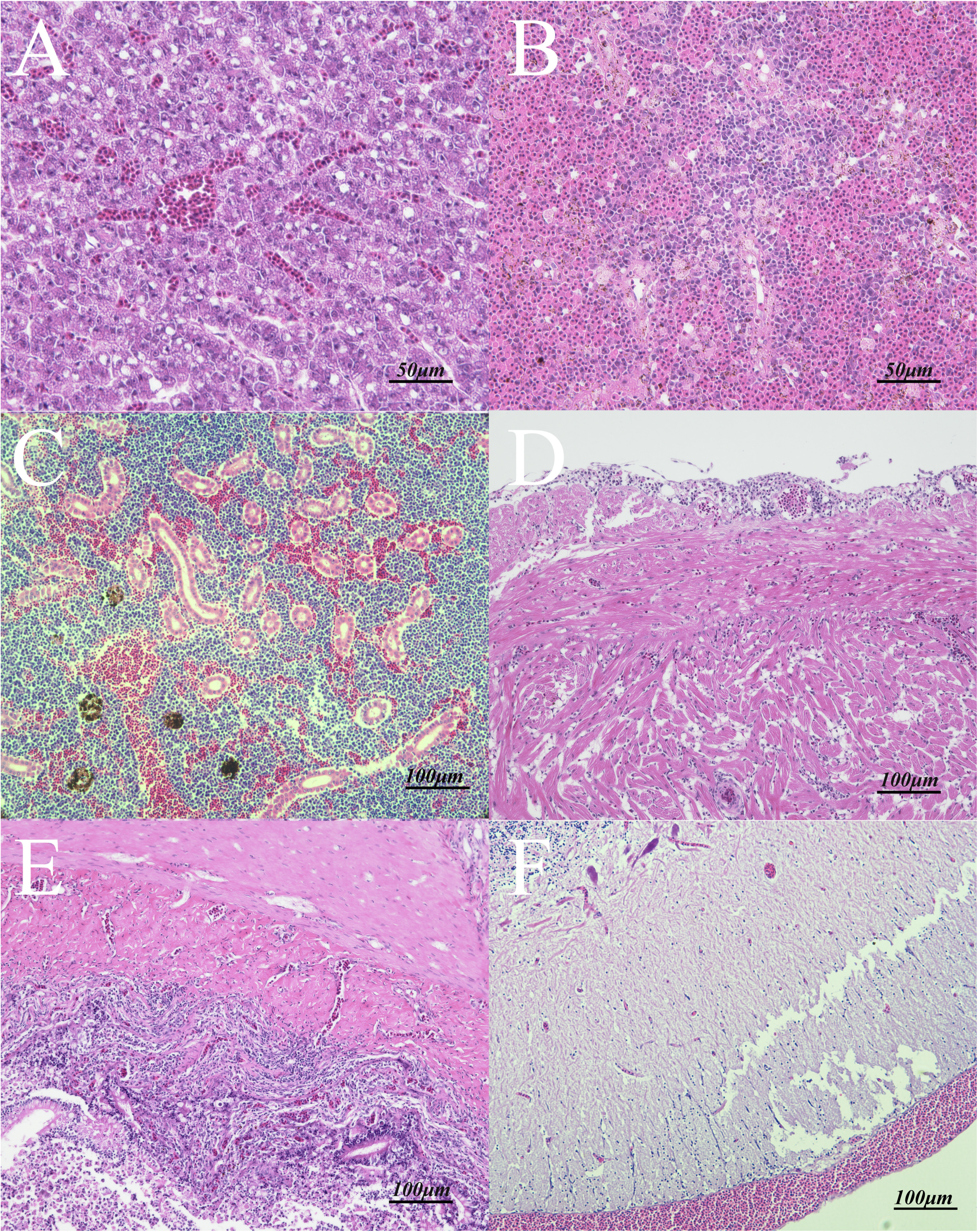
Histological analysis of the naturally infected golden pompano. (A) liver showing hepatocyte swelling and steatosis, along with congestion in hepatic sinuses and blood vessels, (B) splenic tissues exhibiting focal necrosis in the splenic pulp, congestion and hemorrhage in the splenic sinuses and blood vessels, accompanied by the deposition of melanin-macrophage centers. (C) Kidney showing degeneration, necrosis of the renal tubule epithelial cell, focal necrosis, a large amount of inflammatory cell infiltration, and the presence of melanin-macrophage centers in renal hematopoietic tissue. (D) Heart showing epicardial layer was thickened with inflammatory cell infiltration demonstrating the occurrence of epicarditis. (E) Intestine displaying severe necrosis and exfoliation of the intestinal mucosal epithelial cells, a large number of inflammatory cells infiltration in the mucosal layer, and dilated and congested blood vessels in the membrana propria and submucosal. (F) Brain showing obvious cerebrovascular congestion.

### Genome sequencing and Genetic phylogenetic tree analyses

After sequencing, the complete genome of the CAPRV2023 strain from diseased golden pompano was found to be composed of a negative-stranded ssRNA with a size of 11665 nt (GenBank accession no. PP050495). The sequence were predicted to encode N, P, M, G, and L proteins in the order of 3′-N-P-M-G-L-5′ (Figure 7A), which is consistent with the characteristic features of the genus *Novirhabdovirus*. The BLAST analysis showed that the similarity between CAPRV2023 and CAPRV1988 was determined to be 81.32%, whereas the homologies between CAPRV2023 and IHNV, as well as VHSV, were found to be 50.17% and 54.51%, respectively. The phylogenetic tree constructed based on the full genome using the maximum likelihood method in MEGA X revealed that the CAPRV2023 isolate clustered with the CAPRV1988 isolate closely related to the genus *Novirhabdovirus* (Figure 7B).

**Figure 7.**
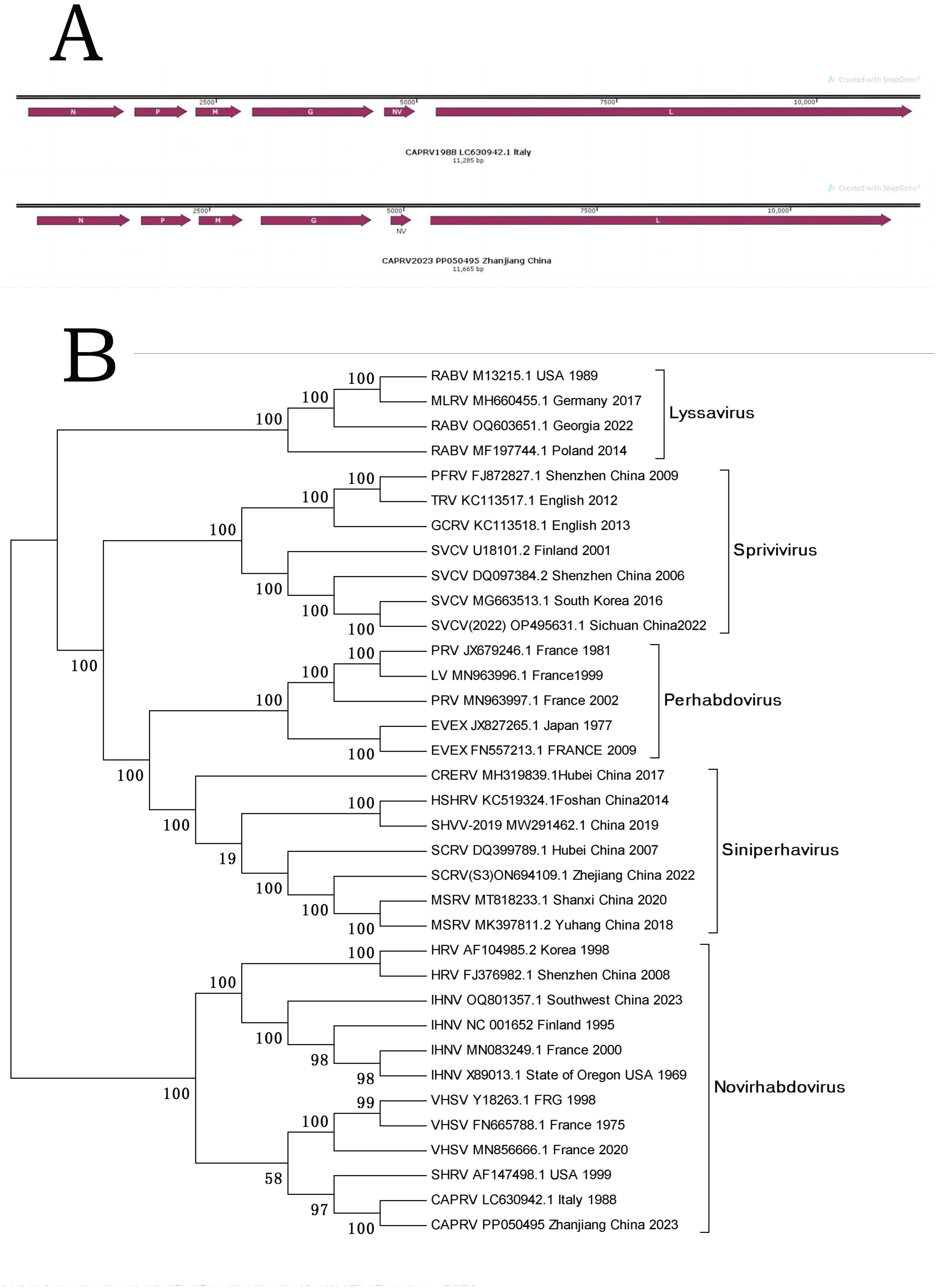
Genome sequencing and genetic phylogenetic tree analyses. (A) Genome map of CAPRV2023. (B) Phylogenetic tree constructed based on the complete genome sequences of CAPRV2023 and other Rhabdomyviridae, using the Clustal W method in MEGA X software with maximum likelihood estimation.

### SNP analysis

The results of SNP analysis are presented in Table 3, with the reference sequence identified as LC630942.1.

**Table 3.**
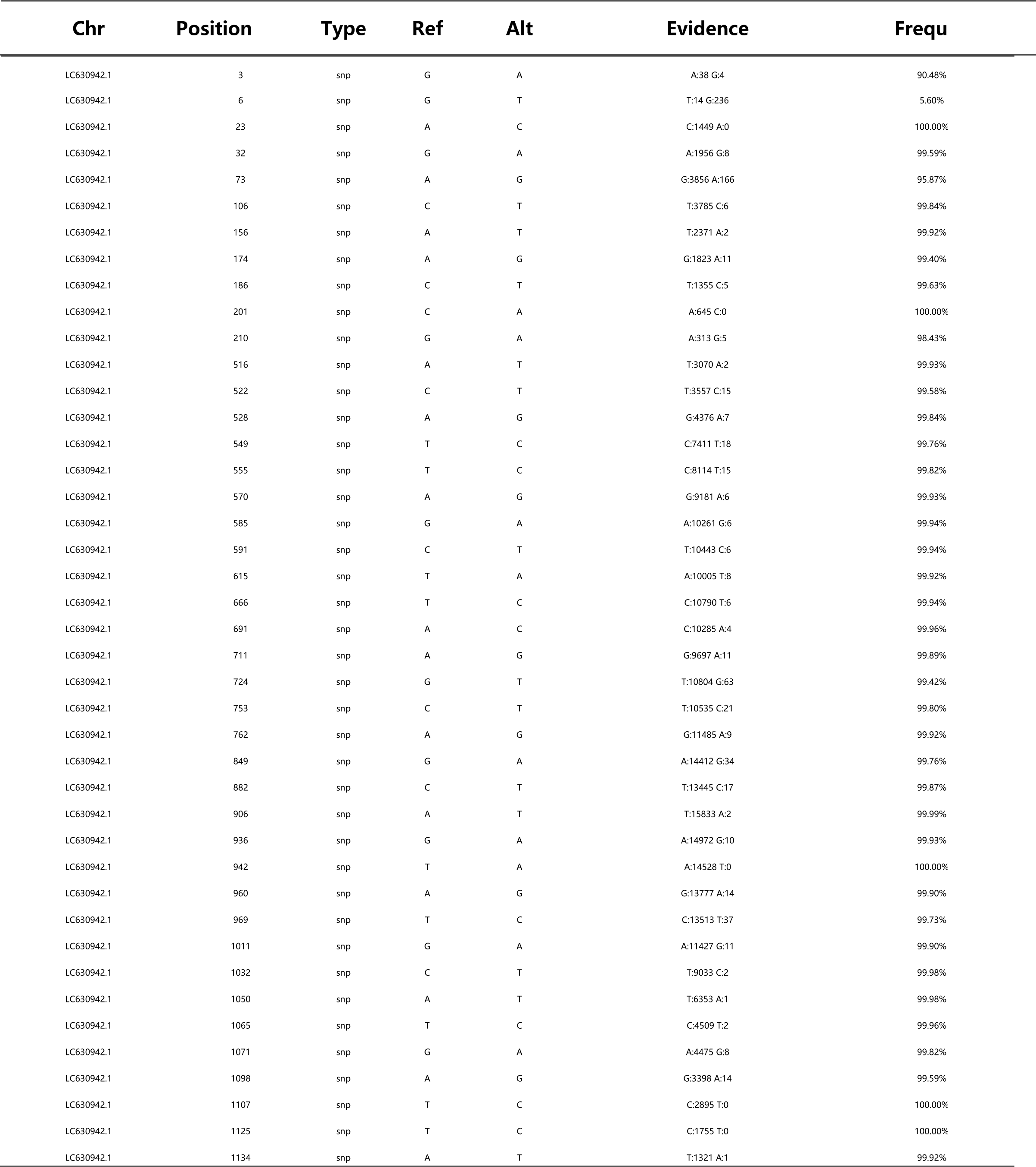
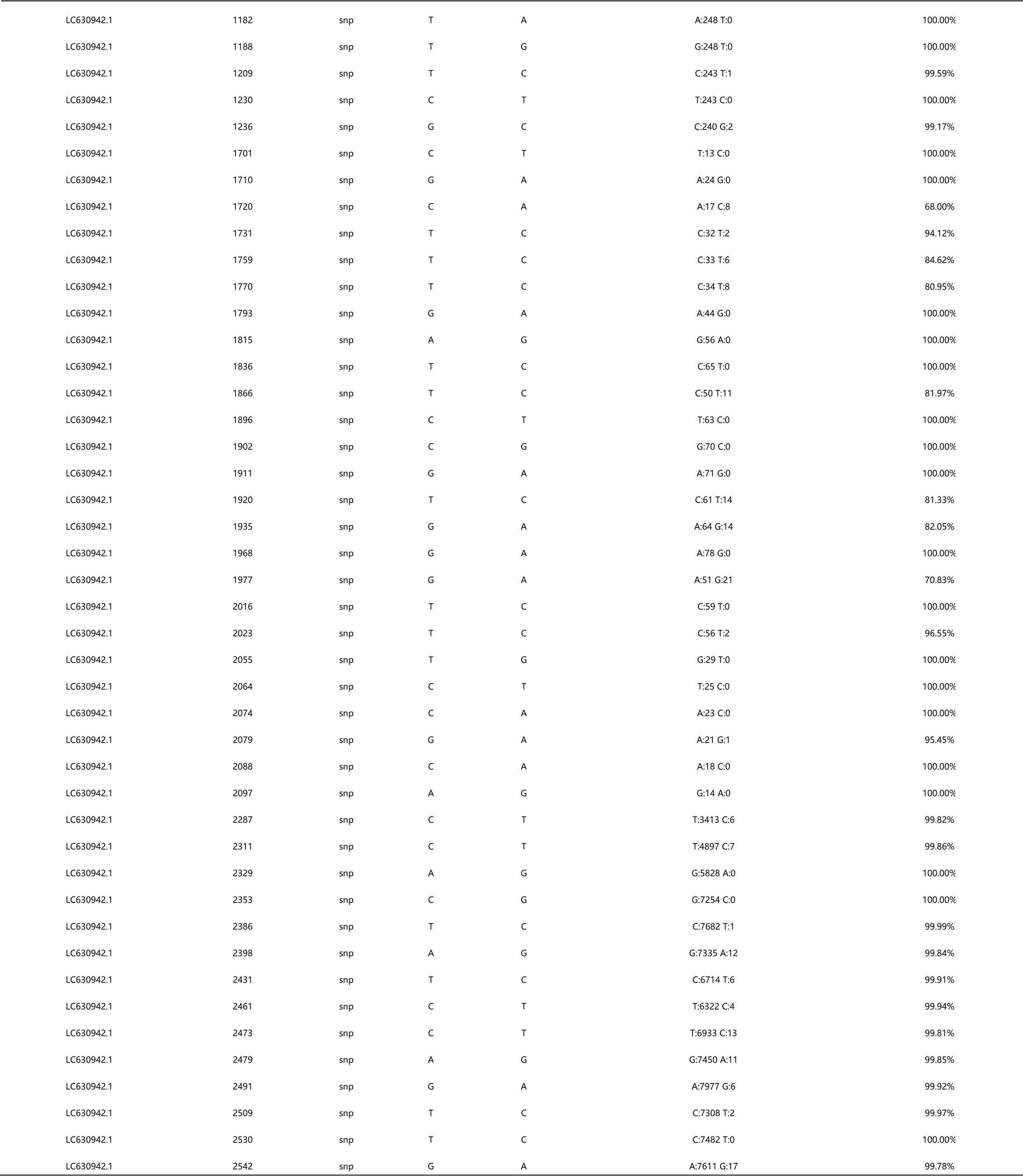
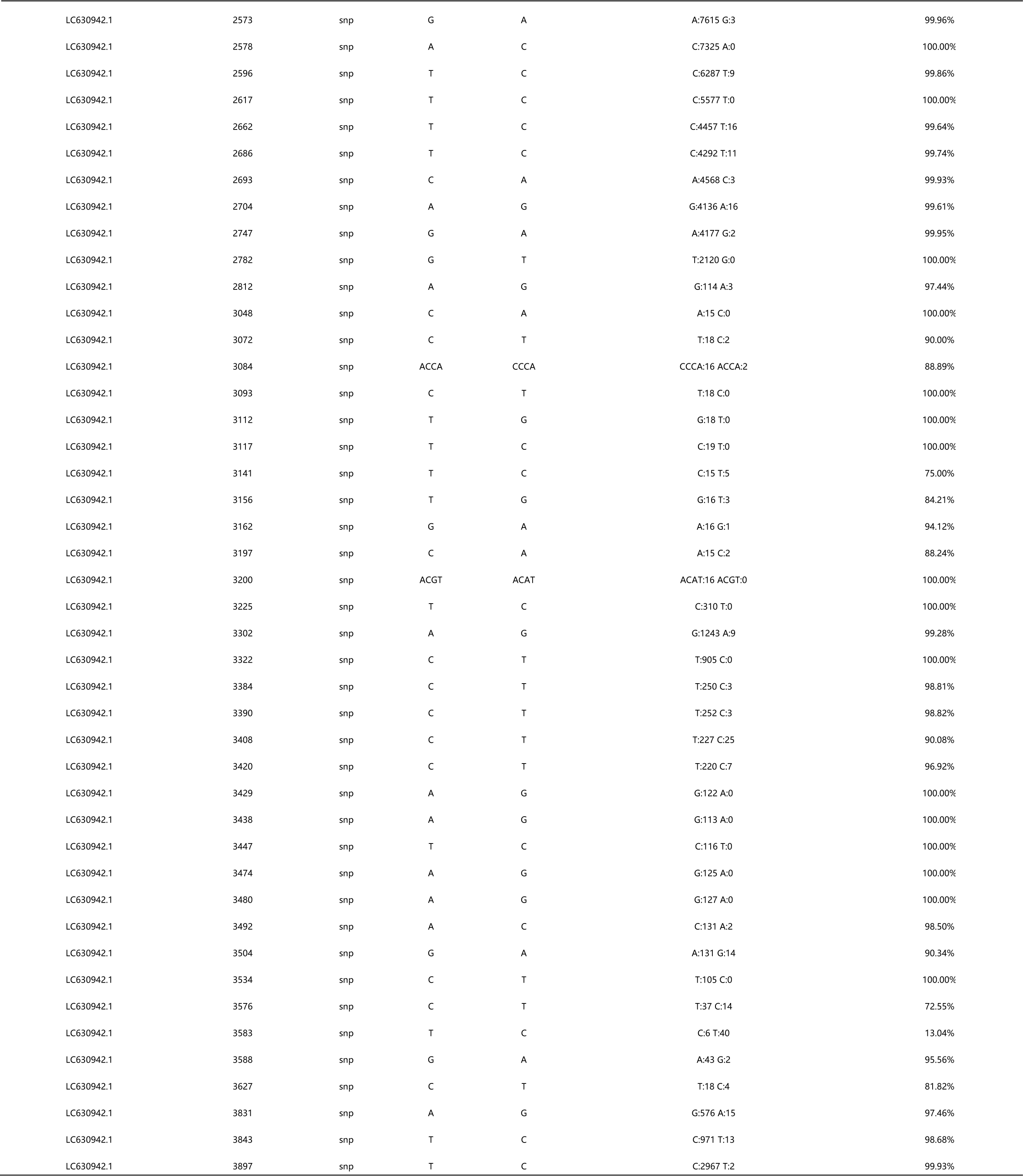
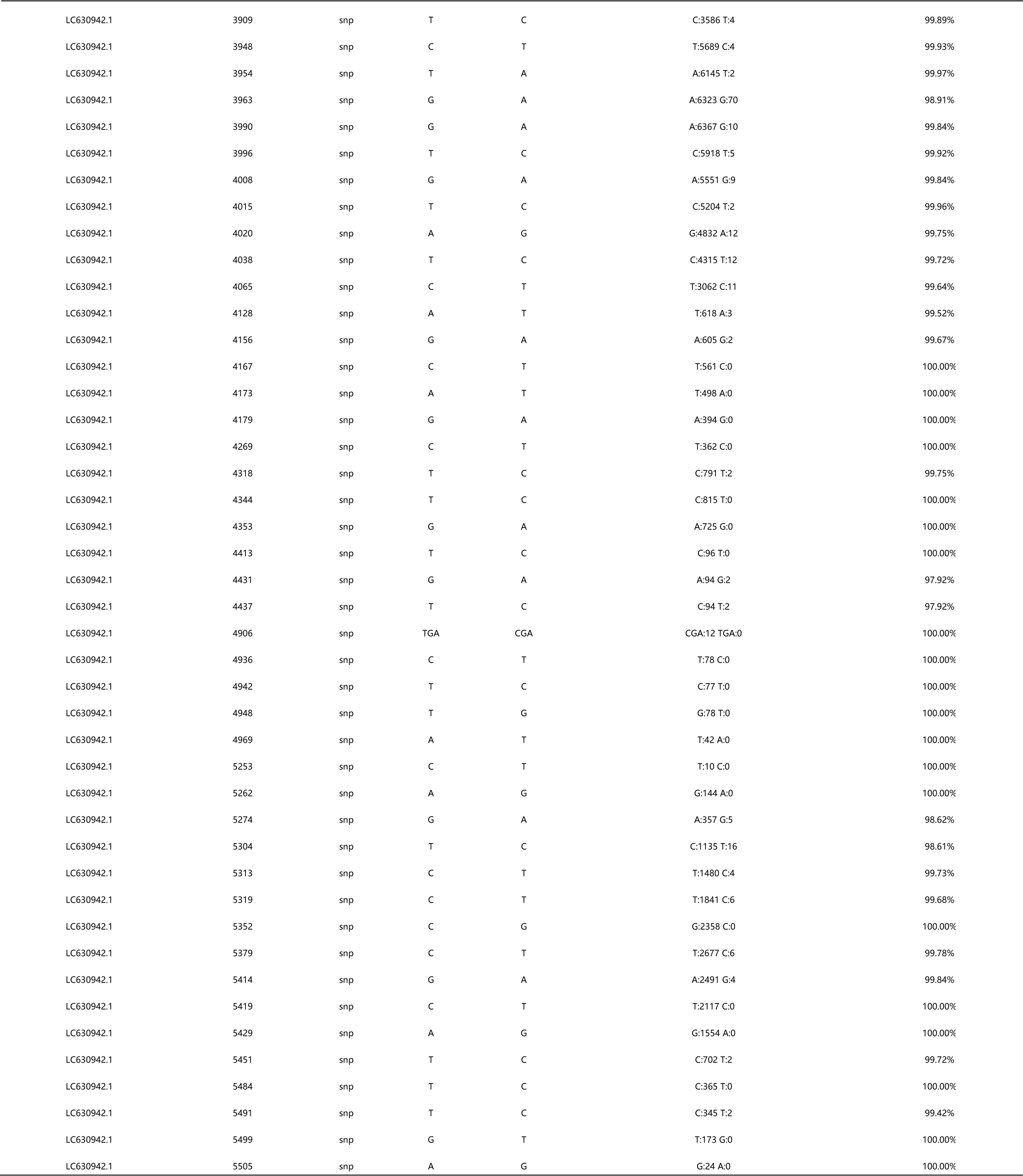
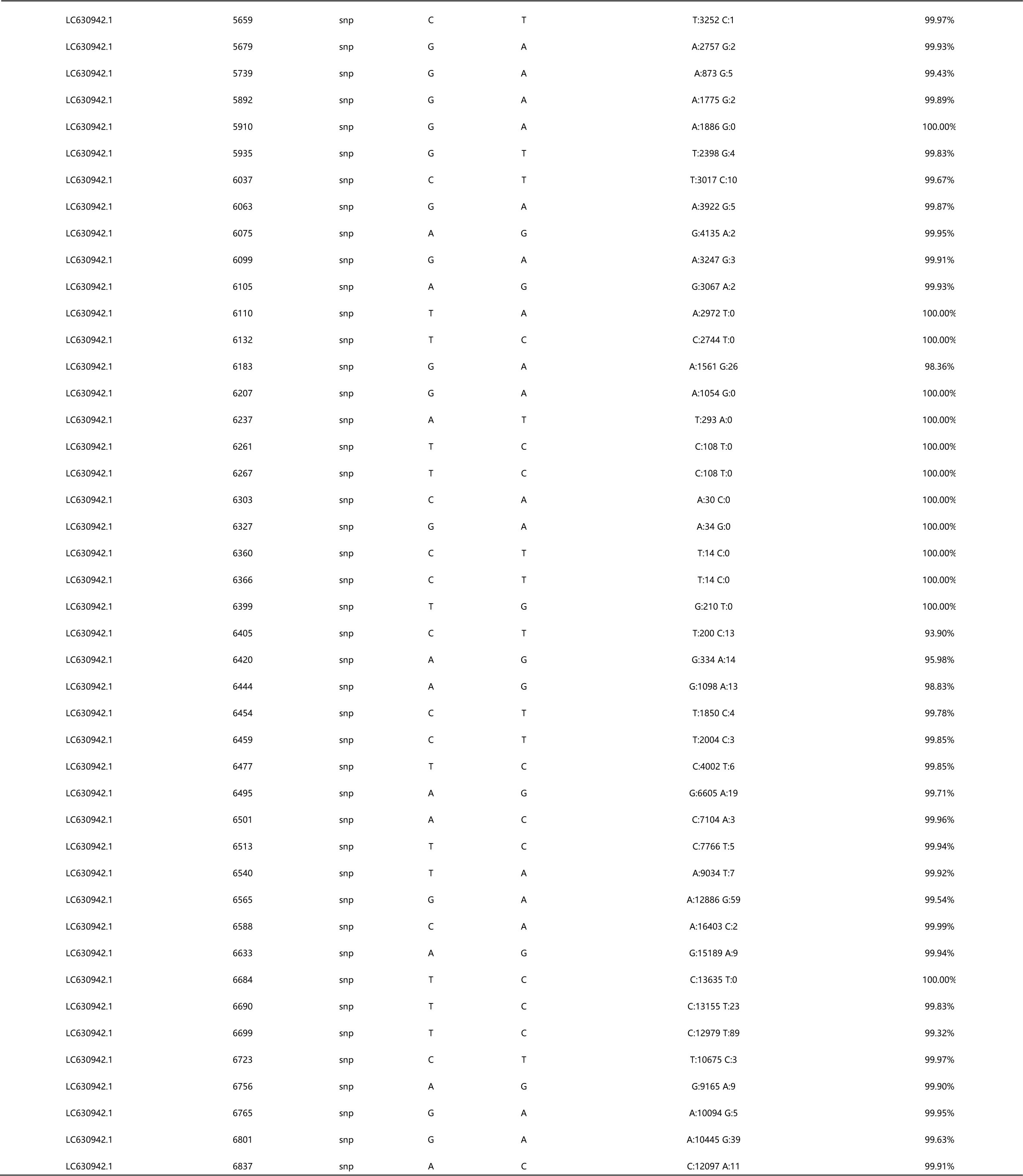
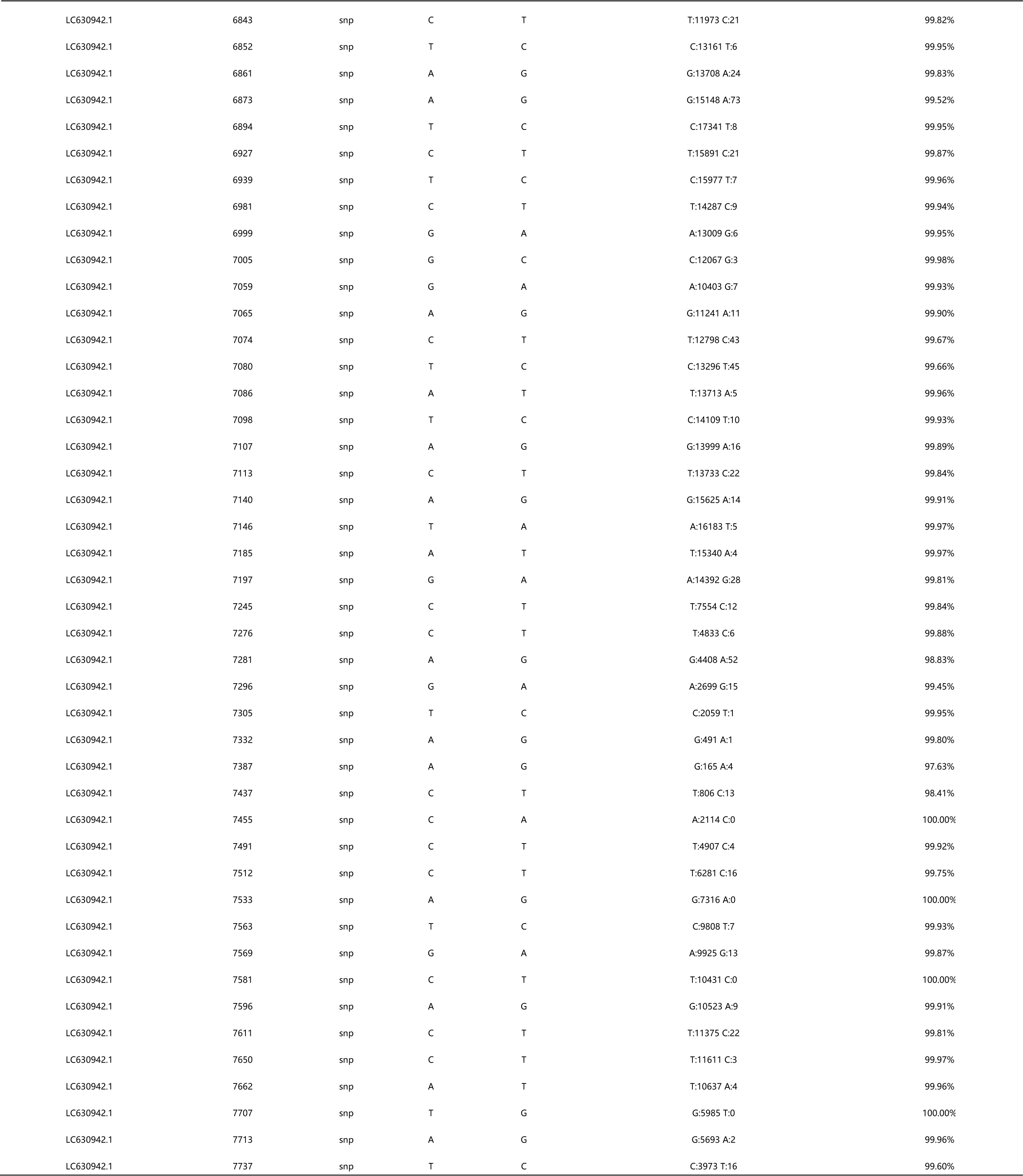
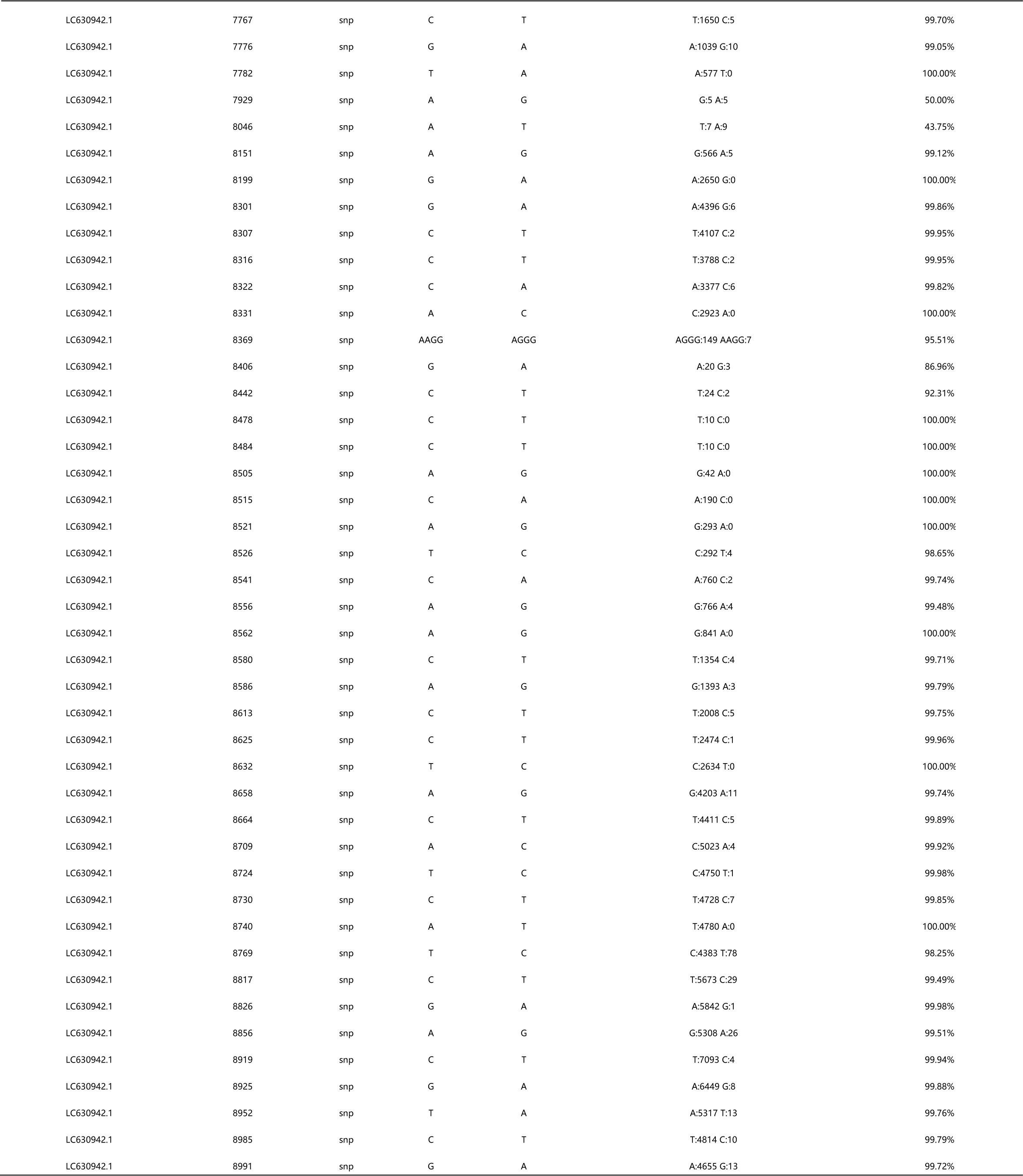
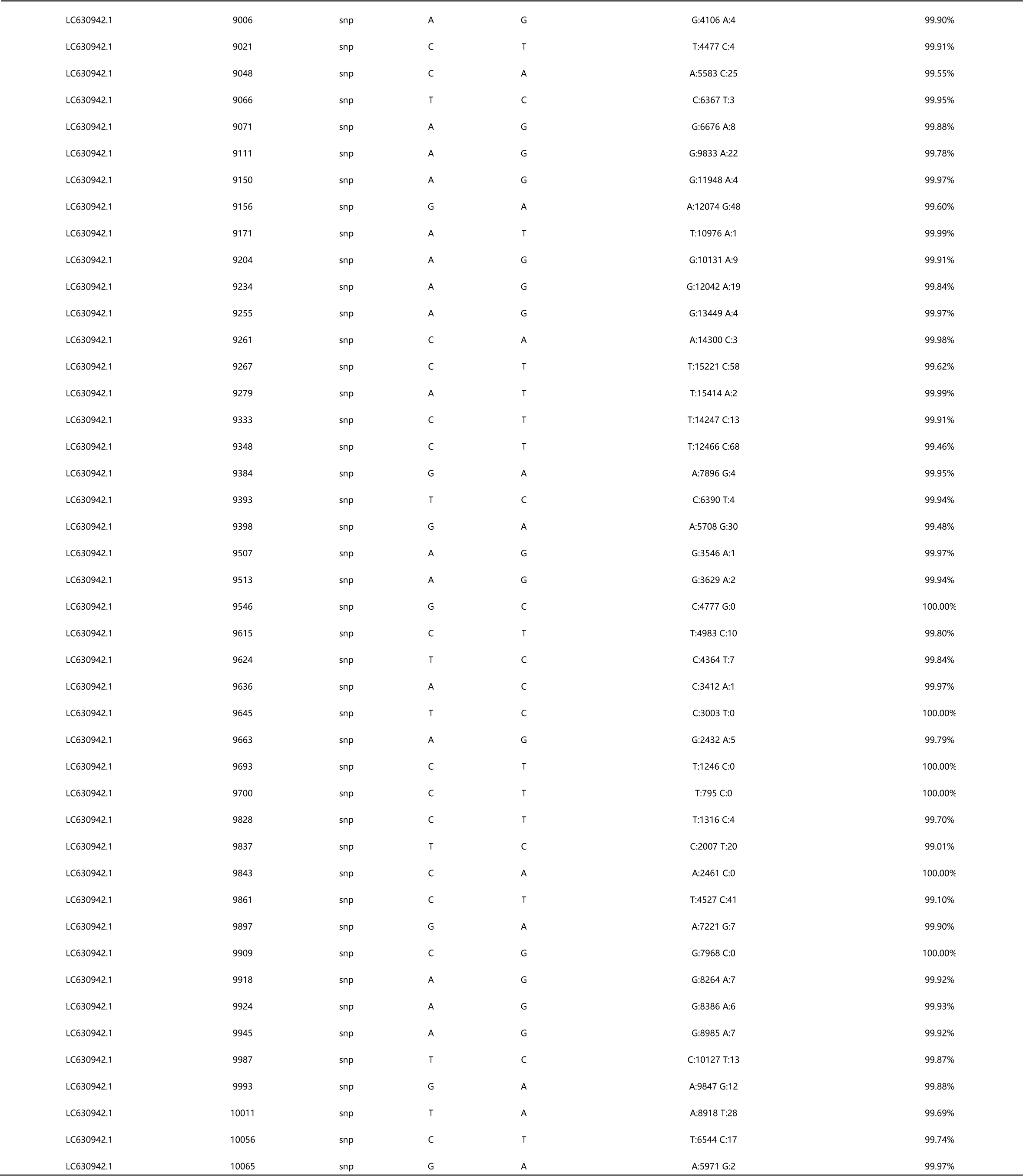
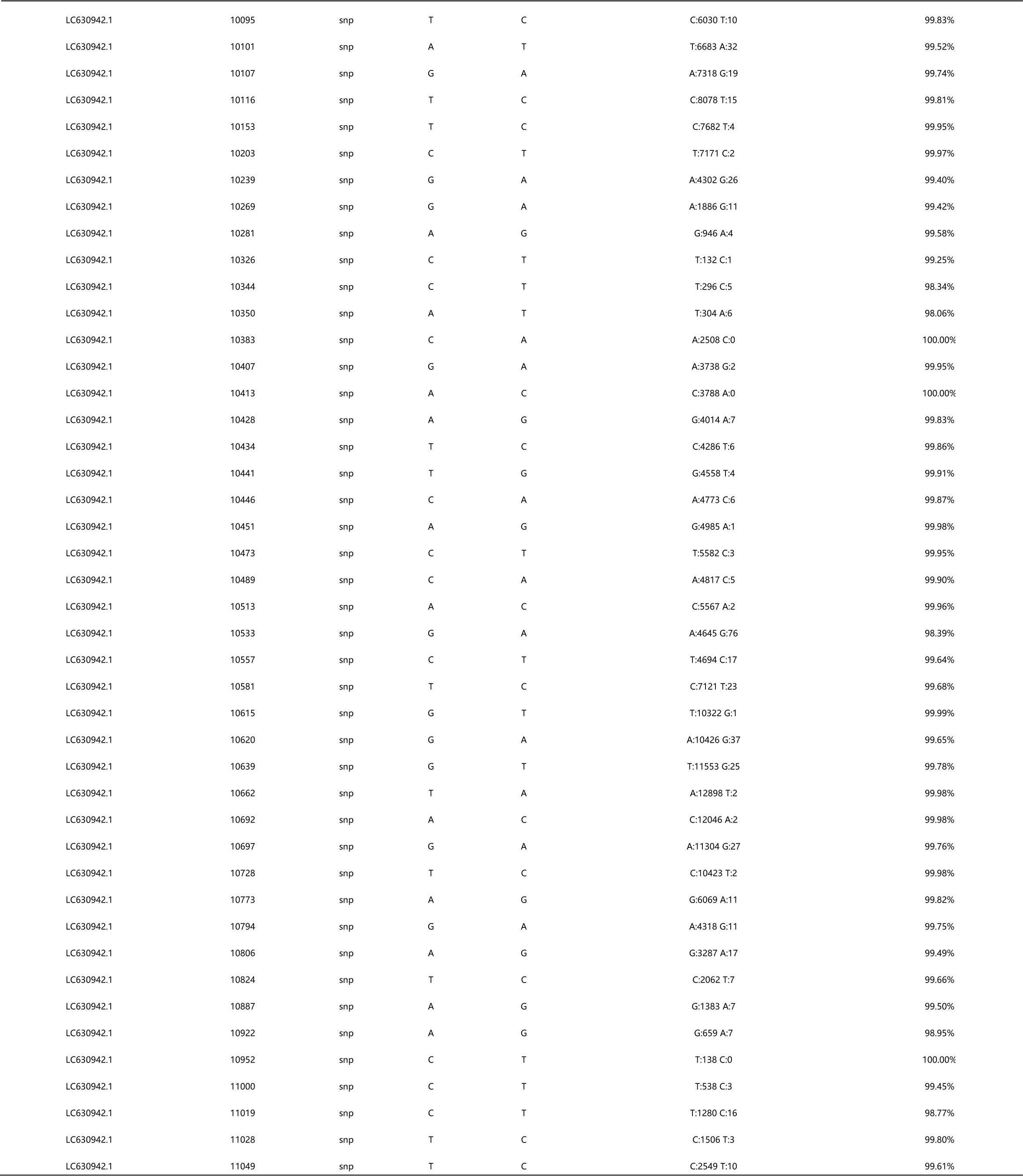
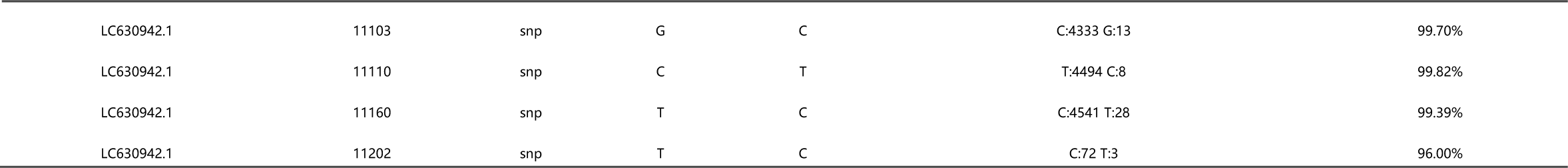
The results of single nucleotide polymorphism (SNP) analysis in CAPRV2023 are consistent with those obtained in CAPRV1988.

### Biological characteristics of Virus

FHM and EPC cell lines were susceptible to the virus isolate CARV-2023. The viral titer in FHM cells (TCID_50_=10^7.33^/0.1 mL) was higher than that in EPC cells (TCID_50_=10^6.28^/0.1 mL) infected with third-passage CAPRV2023.

To determine the effect of temperature on the isolate CAPRV2023 infection, FHM cells infected with CAPRV2023 were cultured at 23, 28, and 33°C, and CPE was also observed at these temperatures, as shown in Figure 8. Remarkably, significant CPE was observed after incubation at 33°C for 8 h, 28°C for 12 h, and 23°C until day 14. After 3 days of infection at 28 and 33°C, almost all cells died. In contrast, only a small number of cells died even after 7 days of infection at 23°C.

**Figure 8.**
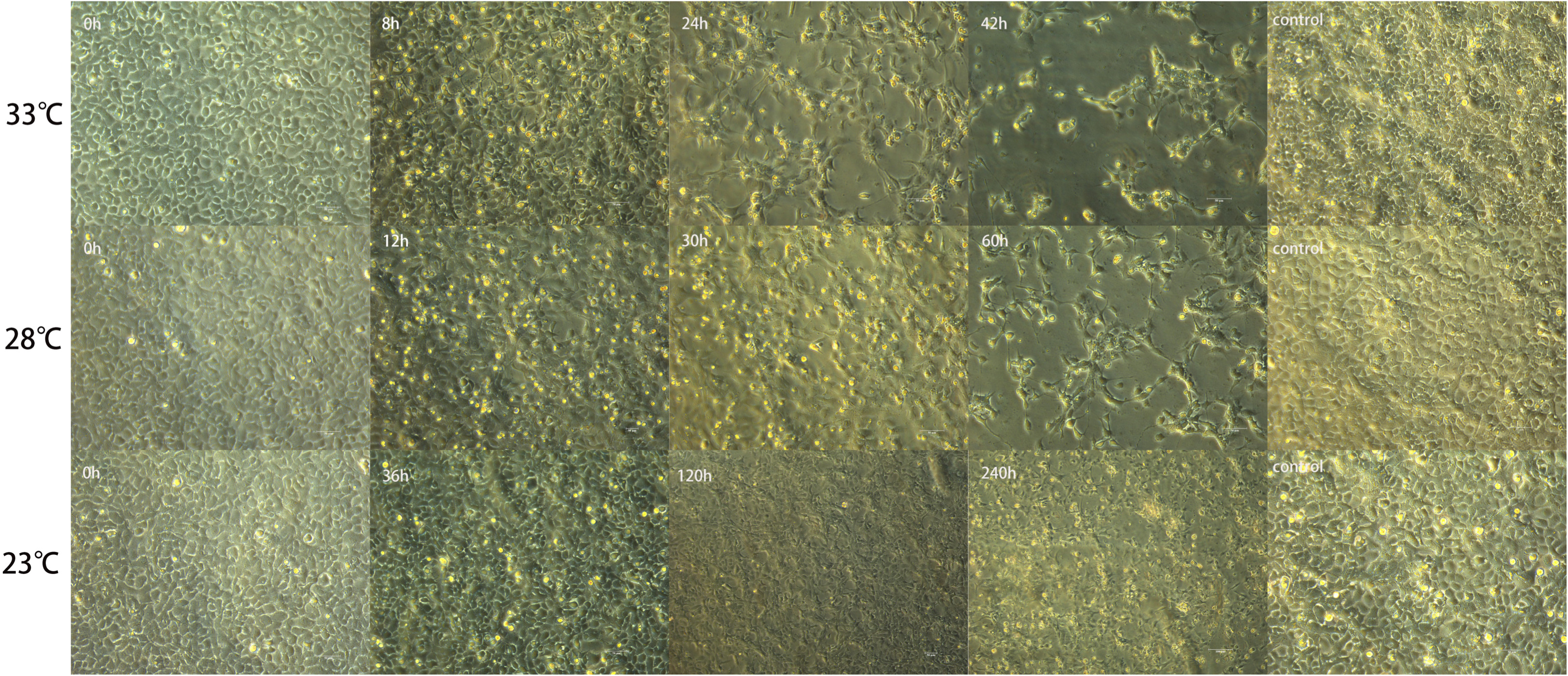
CPE induced by CAPRV2023 at different temperatures.

To further detect replication of CAPRV2023 at different temperatures, FHM cells infected with CAPRV2023 were used to determine the growth kinetics of CAPRV2023 at 23, 28, and 33°C. As shown in Figure 9, the growth kinetic curves of the isolate CAPRV2023 at 28 and 33°C were similar, and the virus titer peaked on days 2 and 3 after incubation and then began to decline. The growth at 23 and 18°C increased slowly (Figure 9). The evidence of viral replication was observed at 10°C. However, the process exhibited an exceptionally slow pace and did not result in significant accumulation during the observation period. (Figure 9).

**Figure 9.**
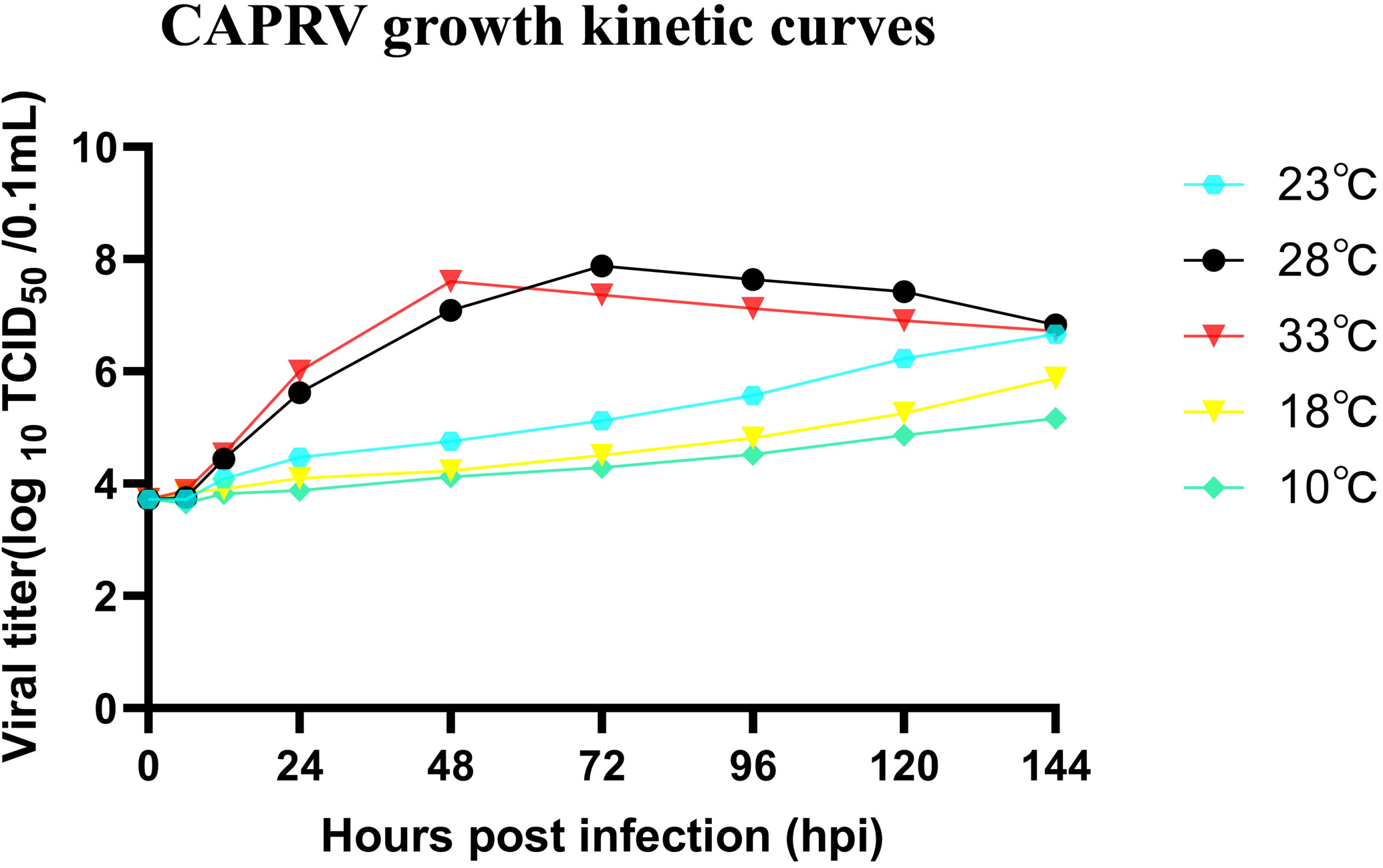
Growth kinetic curves of CAPRV in FHM cells.

To examine the inactivation of CAPRV2023 under UV light, the viral titer was determined after UV irradiation at different time points. As shown in Figure 10, the viral titer decreased rapidly and was reduced by approximately 6 log10 after 45 s of UV irradiation. Further, CAPRV was inactivated at 55°C for 10 min and survived slightly longer at 50 °C but retained some infectivity even after 40 days at 4°C. (Table 2).

**Figure 10.**
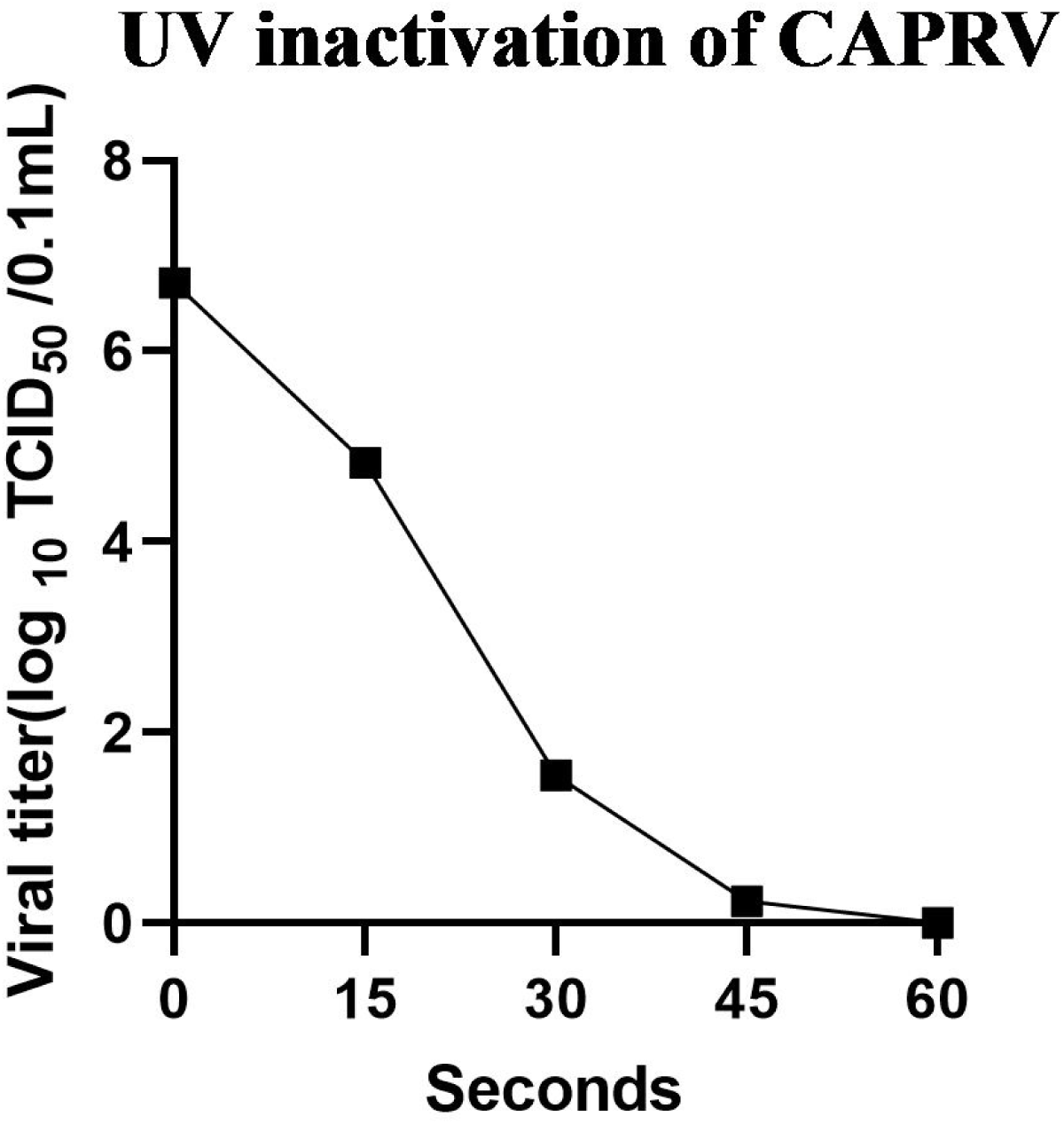
UV irradiation inactivation of CAPRV2023.

The isolate CAPRV exhibited a shorter duration at elevated temperatures and could survival in natural seawater from 21 days at 18°C to 7 days at 33°C. The survival time of CAPRV2023 was slightly shorter in seawater than in ddH_2_O (Figure 11).

**Figure 11.**
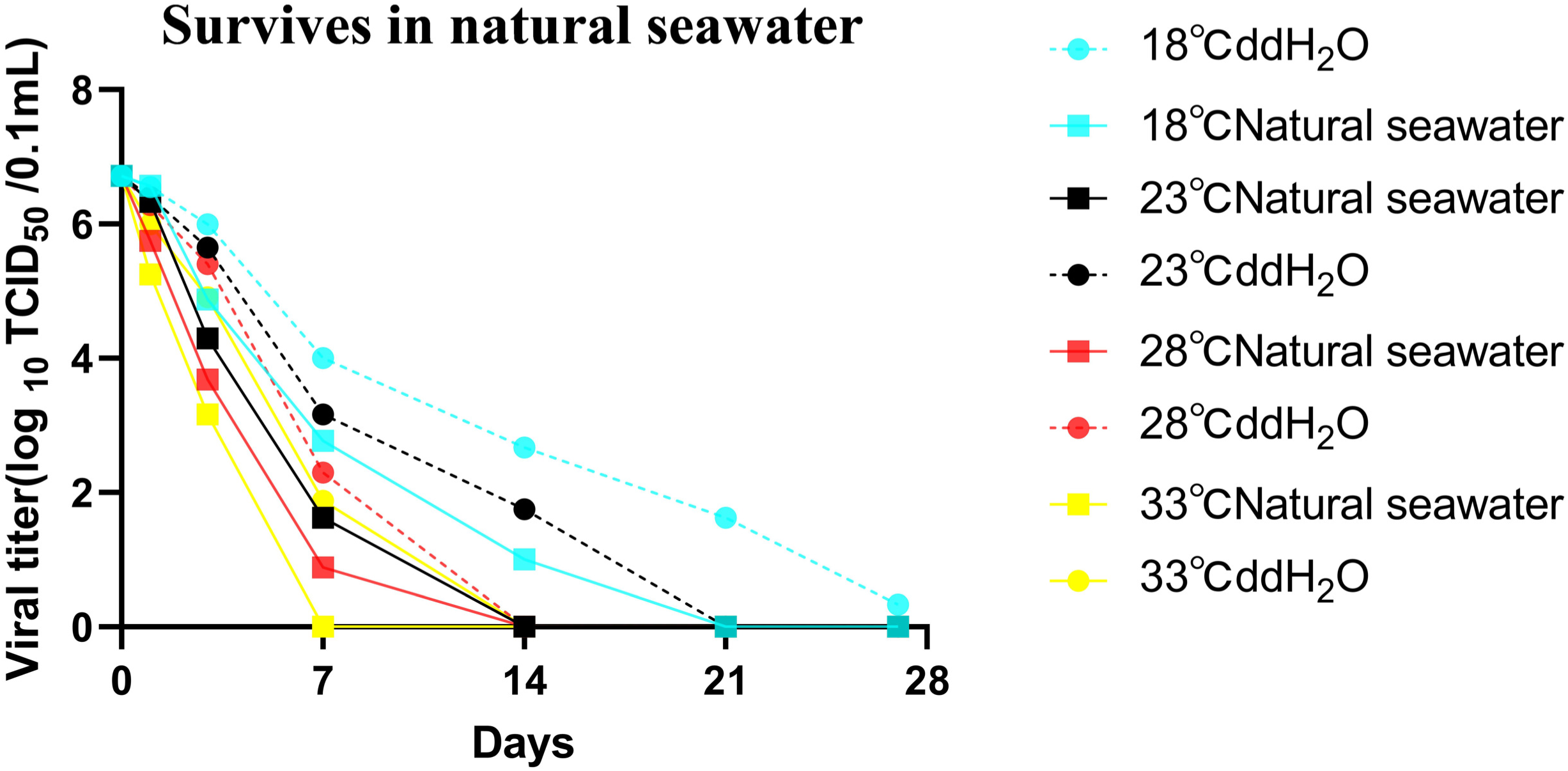
CAPRV2023 survives in natural seawater.

### Fluorescence in situ hybridization

The presence of CAPRV in tissues from naturally infected golden pompanos was detected using fluorescence in situ hybridization. Spleen tissue from a healthy golden pompano was used as a negative control. As shown in Figure 12, the fluorescence signal was the highest in the spleen, followed by the liver, muscles, gill, and kidney and was relatively low in the heart and brain. No positive signals were observed in the spleen tissue of the negative control.

**Figure 12.**
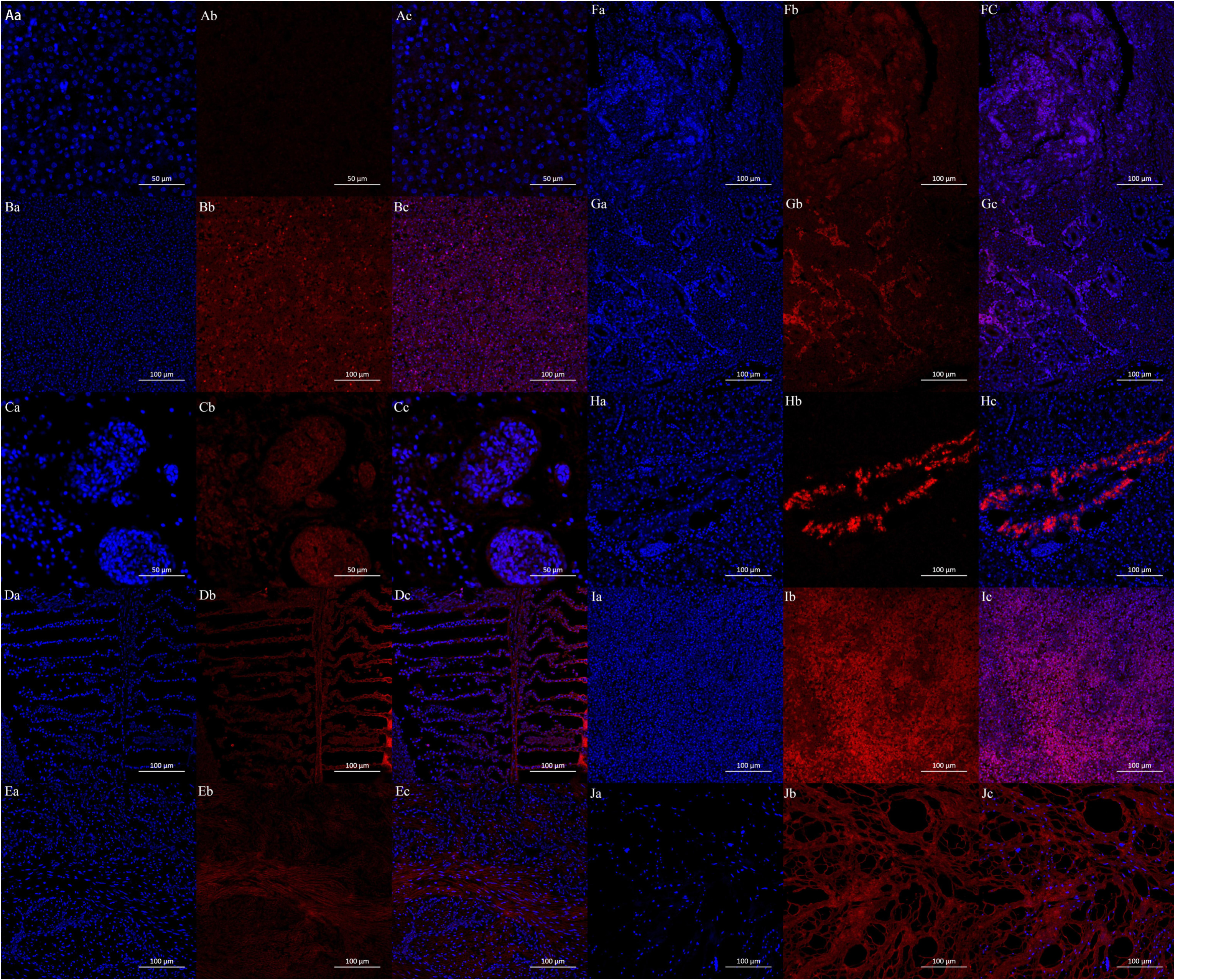
The in situ fluorescence hybridization was performed on each tissue of CAPRV-infected Golden pompano, where the presence of viral proteins is indicated by red fluorescence while the nuclei are stained blue. A-J represent the negative control, positive control, brain, gills, heart, intestine, kidney, liver, spleen and muscle respectively. The intensity of red fluorescence can serve as an indicator for the quantity of viral protein present.

### Experimental infection

Mortality of golden pompano exposed to the isolate CAPRV2023 began from the first day post-infection (dpi), and deaths occurred between 2 and 5 dpi. Fish in the virus-infected groups showed clinical signs of lethargy, anorexia, and whirling movements in water. The cumulative mortality during 3–5 days in the injection infection and immersion injection groups was 100% and 90%, respectively (Figure 13) no morbidity or mortality was observed in the control group. The same viral isolate was re-isolated by cell culture in FHM cells from moribund and dead fish and confirmed using RT-PCR and sequencing (data not shown).

**Figure 13.**
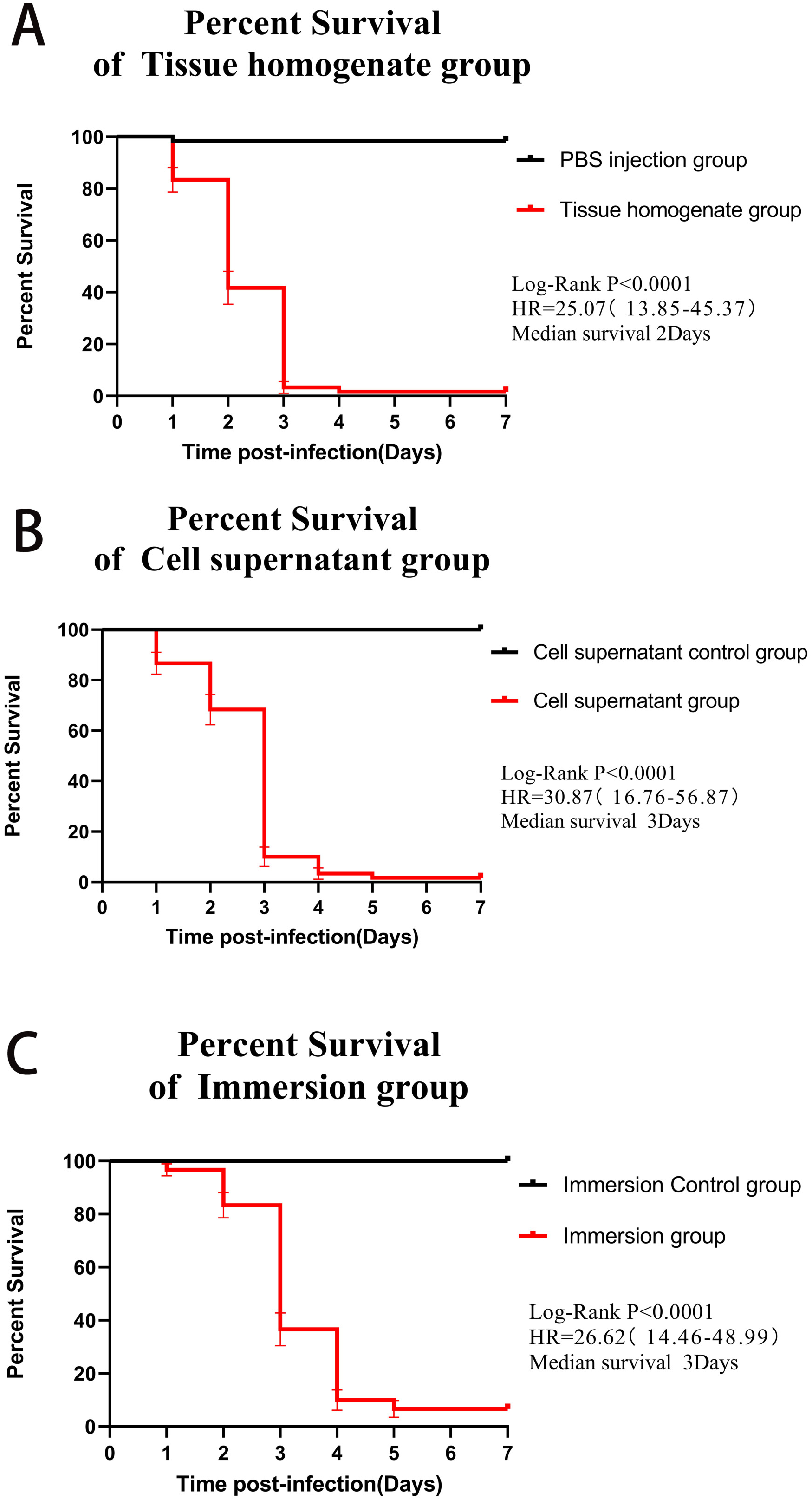
The experimental investigation of CAPRV infection in Golden pompano.

## Discussion

CAPRV was initially isolated from Capione *Salmo trutta* in Italy in 1988 [28], which resulted in high mortality among capiones. Since then, there have been no reports of fish infections caused by this virus. Recently, based on full genome sequencing and serological methods, CARPV was classified as a novel species of *Novirhabdovirus* genus, which differs from other species [29,30]. In the present study, a CAPRV2023 strain was successfully isolated from diseased golden pompanos cultured in offshore cages in China using FHM and EPC cells and detected using nested RT-PCR. The causative agent was confirmed to be a novel CAPRV strain based on morphological analysis, whole-genome sequencing, and artificial infection. To the best of our knowledge, this is the first study to report a natural infection caused by CAPRV in fish in Asian countries. Further, this finding strongly suggests that the golden pompano has emerged as a novel target for CAPRV and has been added to the list of susceptible hosts for CAPRV.

The pathogenicity of rhabdoviruses is usually influenced by temperature, host species, and age [31,32]. Boovo et al. [28] demonstrated that high mortality in Carpio fry exposed to CAPRV at 10 or 15°C. Unexpectedly, mortality was also observed in the control group, probably owing to nutritional deficiencies and inadequate management. The mortality in rainbow trout fry infected with CAPRV was only 12% in the same trial [28]. Severe epizootics caused by CAPRV infections have been observed in offshore cage-cultured golden pompanos in China. CAPRV isolates cause severe disease with high mortality (100%) via the bath route at 28°C. Mortality began earlier in golden pompano (–2–3 dpi) than in Carpiones and rainbow trout (–4–5 dpi). Further, moribund fish showed petechial hemorrhages on the upper and lower jaws, isthmus, and body surface. Experimental infections and findings from natural outbreaks demonstrated that the CAPRV2023 train is highly pathogenic to golden pompano.

The occurrence of infectious diseases in aquatic animals usually results from interactions between the host and the pathogen as well as environmental factors. The correlation between the initiation of certain viruses and environmental factors has been reported in several studies [33,34]. Among these environmental factors, temperature is the key parameter for viral proliferation and pathogenicity because outbreaks of viral diseases in fish are typically temperature dependent [35–45]. For example, IHN outbreaks in salmonid fish usually occur at water temperatures of 8–14°C and have not been observed above 15°C, whereas IHNV replicated most rapidly at 15°C [35,36].

Viral hemorrhagic septicemia (VHS) in Japanese flounder (*Paralichthys olivaceus*), occurs at water temperatures of 8–15°C, whereas the optimum temperature for VHSV replication in vitro is 20 °C[37]. Here, CAPRV could replicate and proliferate over a wider range of water temperatures (23–33°C) during the summer season (August to October). Our results showed that increasing temperature (>23°C) leads to faster viral replication and CPE. This finding suggests that replication efficiency is crucial for CAPRV pathogenicity and may explain the epidemic caused by the CAPR outbreak and high mortality rates experienced during periods of high temperature (>28°C). Further, the epidemic incidence gradually decreased as the temperature decreased and finally disappeared. Therefore, the CAPRV2023 isolate demonstrates thermophilic characteristics (compared to IHNV and VHSV) [36,38] in the same *Novirhabdovirus* genus, which bears a slight resemblance to the SHRV commonly found in tropical regions. The origin of the pandemic remains ambiguous and requires further research.

Although significant CPEs were observed in both FHM and EPC cells following CAPRV infection, the highest viral titers were observed in FHM cells. Therefore, we used FHM cells for the subsequent detection of viral stability. In aquaculture, numerous pathogens can persist in water and spread horizontally, facilitating their transmission to a larger population through water movement. The survival of viruses in natural seawater is influenced by multiple factors. The duration of viral survival in the natural environment was investigated to establish a reliable foundation for epidemiological studies on the virus. Our results demonstrate that CAPRV exhibits enhanced thermal resistance at lower water temperatures, enabling them to survive for longer periods in marine environments. However, even when exposed to the highest recorded water temperature of 33°C in the South China Sea during summer, CAPRV can maintain its infectivity for up to 1–2 weeks. This duration is slightly shorter than that of the SCRV [39] but slightly longer than that of the VHSV [40]. Further, when exposed to CAPRV by immersion in seawater containing a pure culture of the virus, a high mortality rate in the infection group with characteristic clinical signs was observed, suggesting that the virus can be transmitted horizontally. In fact, after the initial discovery of a disease caused by CAPRV in a few offshore aquaculture cages, fish from cages in adjacent waters were also affected by the same disease, further confirming the presence of horizontal transmission. These findings show that the virus could remain infectious in natural marine environments for a long time, posing a potential threat to the culture and disease control of golden pompanos in these regions.

Thermal inactivation at 55°C for 30 min is a commonly used method for virus inactivation. The titer of the virus remained high when SCRV was inactivated at 55°C for 30 min [39]. The temperature and duration required to inactivate the NNV were relatively low [41]. In a study on thermal inactivation, the CAPRV2023 isolate could be completely inactivated following incubation at 55°C for 15 min. These results suggest that viruses attached to aquaculture facilities such as net clothing and fishing nets can be inactivated by prolonged exposure to sunlight [42]. Additionally, UV radiation is another effective way to inactivate viruses in both laboratory and cultural settings. Our results showed that the CAPRV2023 isolate was highly sensitive to UV light and could be completely inactivated following irradiation for 45–60 s, which is in agreement with previous findings on SCRV and VHSV [39,43]. Currently, the possibility of the vertical transmission of CAPRV cannot be ruled out. Therefore, using the UV light to disinfect fertilized eggs and aquaculture water during the larval and juvenile stages is a reliable method for preventing CAPRV infections.

The clinical signs of lethargy, uncoordinated spiral swimming, and hemorrhages around the mouth and on the body surface of the diseased fish in these cases were similar to those observed in other rhabdovirus infections caused by IHNV and VHSV. The pathological lesions observed in the infected fish, including hemorrhages and focal necrosis in the liver, spleen, and kidney, were similar to those reported in farmed Japanese flounder snakehead fish (*Channa striata*) [36,44–47]. In situ hybridization showed that CAPRV2023 was distributed in different tissues, with a higher positive signal intensity in the spleen, liver, and muscle tissues of golden pompano, which was different from that of VHSV in juvenile herring. Using immunohistochemistry, the presence of VHSV was only observed in the kidney and ventricle of the heart of juvenile herring but not in the brain, gills, spleen, or intestinal tissues. Novirhabdovirus distribution in fish could be influenced by virus species, fish species, age, temperature, and other environmental factors [48–50] Notably, the positive reaction of CAPRV2023 in the liver of golden pompano was only observed in blood vessels, which was similar to that in juvenile herring infected with VHSV. Thus, we hypothesized that CRPV, similar to VHSV, targets the vascular endothelium, causing severe petechiae and/or ecchymotic hemorrhage in the skin, muscles, and internal organs. Further studies are required to identify these target cells.

Although CAPRV was discovered in 1988, no rapid detection technology currently exists for the virus. Virus detection relies mainly on cell culture, genome sequencing, and animal experiments, which are both time consuming and labor intensive. Therefore, the development of specific rapid detection techniques for CAPRV is urgently required. We successfully developed a fundamental RT-PCR and nested PCR technique for the rapid detection of CAPRV in the laboratory. The developed method has outstanding specificity and sensitivity in laboratory and clinical tests, providing a powerful tool for the early detection of diseases and epidemiological research.

In conclusion, through cell isolation, infection testing, and histopathological analysis, the causative agent responsible for the golden pompano epidemic was determined to be CAPRV. Further, some biological characteristics of the virus have been characterized. We also developed an ISH assay and a nested PCR for the specific detection of CAPRV. It is imperative to prioritize the epidemiological investigation of the virus, vigorous surveillance programs, research on pathogenic mechanisms, implementation of biosecurity measures, and development of vaccines to mitigate the impact of this emerging viral disease on the golden pompano industry.

## Acknowledgements

This work was funded by the National Key Research and Development Program (No.2022YFD2401204) and Key-Area Research and Development Program of Guangdong Province (No.2021B0202040002).

## Institutional Review Board Statement

All animal experiments were conducted strictly based on the recommendations in the ‘Guide for the Care and Use of Laboratory Animals’set by the National Institutesof Health. All fish experiments were approved by the Guide of the Animal Ethics Committee of Guangdong Provincial Key Laboratory of Aquatic Animal Disease Control and Healthy Culture., approval code: (20230915)002, approval date: 15 September 2023.

